# Dissecting the Roles of Mitochondrial Complex I Intermediate Assembly (MCIA) Complex Factors in the Biogenesis of Complex I

**DOI:** 10.1101/808311

**Authors:** Luke E. Formosa, Linden Muellner-Wong, Boris Reljic, Alice J. Sharpe, Traude H. Beilharz, Michael Lazarou, David A. Stroud, Michael T. Ryan

## Abstract

Mitochondrial Complex I harbors 7 mitochondrial and 38 nuclear-encoded subunits. Its biogenesis requires the assembly and integration of distinct intermediate modules, mediated by numerous assembly factors. The Mitochondrial Complex I Intermediate Assembly (MCIA) complex, containing assembly factors NDUFAF1, ECSIT, ACAD9, and TMEM126B, is required for building the intermediate ND2-module. The role of the MCIA complex and the involvement of other proteins in the biogenesis of this module is unclear. Cell knockout studies reveal that while each MCIA component is critical for complex I assembly, a hierarchy of stability exists centred on ACAD9. We also identify TMEM186 and COA1 as *bona fide* components of the MCIA complex with loss of either resulting in in MCIA complex defects and reduced complex I assembly. TMEM186 enriches with newly translated ND3, while COA1 enriches with ND2. Our findings provide new functional insights into the essential nature of the MCIA complex in complex I assembly.

## Introduction

Mitochondrial complex I (NADH:ubiquinone oxidoreductase) is the first component of the oxidative phosphorylation (OXPHOS) system and is required for the generation of the proton motive force driving ATP production and normal mitochondrial function (Hirst, 2013; Sazanov, 2015). In humans, complex I is composed of 45 subunits encoded by both the nuclear and mitochondrial genomes and requires at least 15 assembly factors for its biogenesis (Formosa et al., 2018; Sánchez-Caballero et al., 2016a; Zhu et al., 2016). Of the 45 subunits, 14 core subunits are required for substrate oxidation, electron transport and proton translocation and are highly conserved from bacteria to mammals. Recent high resolution cryo-EM structural studies of mammalian complex I have shown that the remaining 30 accessory subunits form a proteinaceous cage that surround the core complex I subunits (Fiedorczuk et al., 2016; Zhu et al., 2016), and most are required for assembly, stability and hence activity of the enzyme (Stroud et al., 2016). Complex I further associates with respiratory complexes III and IV to form the respiratory supercomplex, the function of which is still under debate (Fedor and Hirst, 2018; Guo et al., 2017; Letts et al., 2016; Milenkovic et al., 2017). It has also been demonstrated that loss of complex III reduces the levels of/or impedes assembly of complex I, suggesting a dynamic interplay between the respiratory enzymes during their biogenesis (Acín-Pérez et al., 2004). Indeed, models for the assembly of the supercomplex include partially assembled complex I integrating with free subunits or assembly intermediates of complexes III and IV (Moreno-Lastres et al., 2012), while other evidence supports the formation of the supercomplex from pre-existing, fully assembled enzymes (Acin-Perez et al., 2008; Guerrero-Castillo et al., 2017)

The current model of complex I assembly (Formosa et al., 2018; Sánchez-Caballero et al., 2016a) involves the synthesis and inner membrane insertion of seven core mtDNA-encoded subunits – ND1, 2, 3, 4, 4L, 5 and ND6. Insertion appears to take place in a co-translational manner via the insertase OXA1L (Thompson et al., 2018). The remaining subunits - seven core and 30 accessory subunits - are all nuclear-encoded and are imported into mitochondria. The majority are imported into the matrix where they assemble into matrix arm intermediates (Q- and N-modules) while other subunits assemble in specific modules at the membrane with mtDNA-encoded subunits (Dunning et al., 2007; Lazarou et al., 2007; Stroud et al., 2016; Ugalde et al., 2004; Vogel et al., 2007a). In cells with mutations in mtDNA-encoded subunits, a number of assembly intermediates have also been observed that accumulate with both mitochondrial and nuclear encoded subunits (Perales-Clemente et al., 2010). This includes the ND1-module (also referred to as P_P_-a) and ND2-module (P_P_-b) that are part of the proximal membrane arm and the ND4-module (P_D_-a) and ND5-module (P_D_-b) that are part of the distal membrane arm (Guerrero-Castillo et al., 2017; Sánchez-Caballero et al., 2016a). Assembly factors play a prominent role in the biogenesis of complex I and perform functions that include the post-translational modification of subunits (Rhein et al., 2013; Rhein et al., 2016; Zurita Rendon et al., 2014), delivery of cofactors (Sheftel et al., 2009), insertion of proteins into the inner membrane, and stabilization of partially assembled, intermediate modules (Formosa et al., 2018). In cases where complex I subunits are damaged or assembly is compromised, modified, unincorporated, labile or misassembled subunits are degraded by mitochondrial proteases to maintain mitochondrial integrity or reduce levels of reactive oxygen species (Pryde et al., 2016; Puchades et al., 2019; Rendón and Shoubridge, 2012; Stiburek et al., 2012).

Many mutations in genes encoding assembly factors have been identified in cases of complex I deficiency and mitochondrial disease, highlighting an important role of these proteins in cell and tissue function (Fiedorczuk and Sazanov, 2018; Formosa et al., 2018). In this study, we focus on the role of the Mitochondrial Complex I Intermediate Assembly (MCIA) complex, an inner membrane machine composed of numerous assembly factors whose function is poorly understood, but converges on the biogenesis of the ND2-module. The MCIA complex is composed of core subunits NDUFAF1 (Vogel et al., 2005), ECSIT (Vogel et al., 2007b), ACAD9 (Nouws et al., 2010) and TMEM126B (Heide et al., 2012). The assembly factor TIMMDC1 has also been identified in association with the MCIA complex and stalled complex I intermediate (Andrews et al., 2013; Guarani et al., 2014). Pathogenic mutations have also been identified in NDUFAF1 (Dunning et al., 2007; Fassone et al., 2011), ACAD9 (Haack et al., 2010; Nouws et al., 2010), TMEM126B (Alston et al., 2016; Sánchez-Caballero et al., 2016b) and TIMMDC1 (Kremer et al., 2017). In addition, complexome analysis – correlative analysis of proteins that migrate together on blue-native gels – revealed that TMEM186 and COA1 co-migrate with complex I assembly intermediates that contain known MCIA complex components (Guerrero-Castillo et al., 2017). As yet, a direct role for TMEM186 and COA1 in complex I assembly has not been experimentally validated.

Here we characterise the MCIA complex and identify the known core subunits being critical for complex I assembly with a dynamic interplay between the stability of subunits, while TMEM186 and COA1 serve important, yet more peripheral roles in assembly of the enzyme.

## Results

### Loss of complex I accessory subunits perturbs steady-state assembly factor complexes

Complex I is assembled via the joining of different modules. In an elegant study, Nijtmans and colleagues profiled the *de novo* assembly of complex I using Blue Native (BN)-PAGE and quantitative mass-spectrometry, and uncovered at least 10 distinct intermediate complexes containing various assembly factors (Guerrero-Castillo et al., 2017). This included the MCIA and TIMMDC1 modules forming an intermediate containing the Q/ND1 and ND2 modules (designated Q/P_P_). The MCIA-ND2 module was also found in an independent ND4 assembly (P_P_-b/P_D_-a). However, under homeostatic conditions where a population of complex I and intermediates is always present, the MCIA and TIMMDC1 complexes resolve as a number of more defined complexes on BN-PAGE (**Fig. 1**). ACAD9-containing complexes from control mitochondria consist of a major species at ∼450kDa (**Fig. 1A, lanes 1 and 32, marked with #**) and a larger doublet of ∼680kDa and ∼720kDa (**Fig. 1A, lanes 1 and 32, marked with \***). TIMMDC1 is found in steady state complexes of ∼400 kDa, 440 kDa (**Fig. 1B, lanes 1 and 32, marked with †**) plus two larger complexes similar to that seen for ACAD9 (**Fig. 1B, lanes 1 and 32, \***).

**Figure 1:**
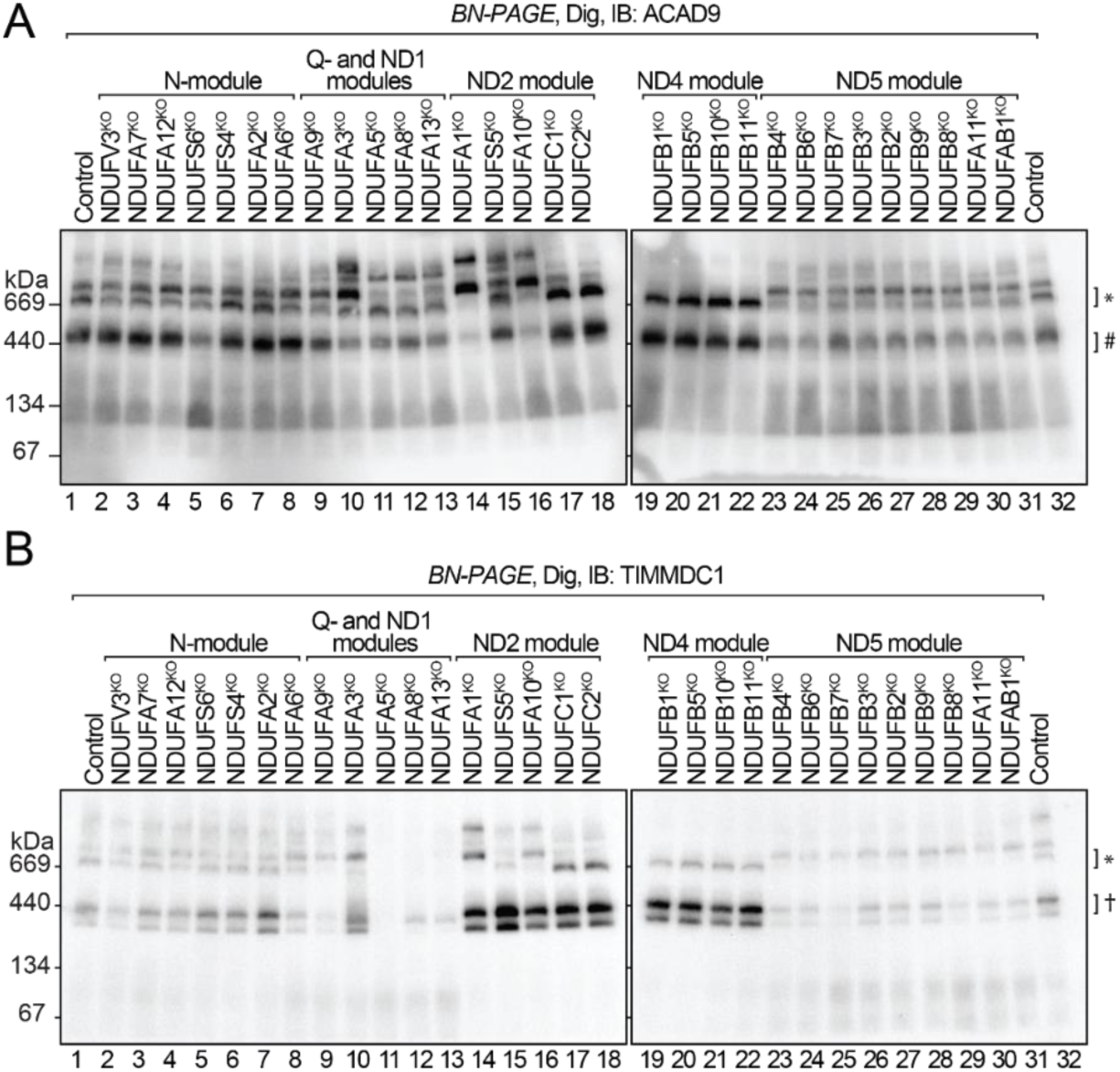
Complex I assembly intermediates are disrupted in accessory subunit KO mitochondria. Mitochondria from indicated cell lines were solubilized in digitonin and analysed by BN-PAGE and immunoblotting (IB) for **(A)** ACAD9 and **(B)** TIMMDC1. #, ACAD9 ∼450kDa complex. †, TIMMDC1 ∼400 and 440kDa complexes. *, ∼680 and 720kDa complexes.

We first sought to determine how these complexes were affected in mitochondria lacking individual complex I subunits belonging to defined modules that we have previously characterised (Stroud et al., 2016). Loss of subunits from the NADH-dehydrogenase (N-) module did not affect the appearance of either ACAD9 or TIMMDC1 modules (lanes 2-8), supporting earlier studies that N-module assembly occurs at the last steps in assembly (Lazarou et al., 2007; Stroud et al., 2016). Although reduced in levels, the migration of these complexes was also largely unaffected in mitochondria lacking subunits belonging to the distal ND5 module (lanes 23-31). Variations were clearly noticeable in subunit knockouts belonging to the Q, ND1, ND2 and ND4 modules. The absence of the ND4 module subunits led to specific loss of the higher complex for both ACAD9 and TIMMDC1 (lanes 19-22). This suggests that the higher species represents assembly factors associated in an intermediate containing the Q/ND1/ND2/ND4 module (i.e. Q/P_P_/P_D_-a). Furthermore, loss of ND4-module subunits resulted in robust accumulation of the TIMMDC1 doublet (**Fig. 1B, lanes 19-22, labelled with †**), with similar changes observed when ND2 module subunits are missing (**Fig. 1B, lanes 14-18**). Changes to the higher complexes containing TIMMDC1 in mitochondria lacking ND2-module subunits mirrored those observed for ACAD9 (except for NDUFS5^KO^ mitochondria), suggesting that most of these complexes represent associations between assembly factors, the ND1-module and crippled ND2-modules. Finally, in mitochondria lacking some of the Q- and ND1-module subunits, ACAD9-containing complexes were perturbed with loss of the accumulation of a higher molecular weight species (**Fig. 1A, lanes 10-13**). TIMMDC1 complexes were also absent, with the exception of NDUFA3^KO^ mitochondria (**Fig. 1B, lanes 9-13**). TIMMDC1 complexes were also absent from NDUFA5^KO^ mitochondria. NDUFA5 is an accessory subunit of the Q-module, suggesting proper formation of the Q-module is a requirement for TIMMDC1 to form the 400kDa and 440kDa complexes. TIMMDC1 complexes were also absent in NDUFA8 and NDUFA13 knockout mitochondria, except for a low abundant species migrating between the 400 and 440kDa complexes. In summary, we conclude that ACAD9 and TIMMDC1 exist in independent lower molecular weight steady state complexes as well as together in higher molecular weight species. Our results also suggest that disruption of any single complex I assembly module can have a large effect on the behaviour of other modules of the assembly pathway, and may not be restricted to the modules directly affected.

### The MCIA complex is crucial for complex I assembly

To investigate the interplay of the assembly factors in more detail, we generated KO cell lines for MCIA genes *ACAD9, ECSIT* and *TMEM126B* and used these together with previously-validated NDUFAF1^KO^ and TIMMDC1^KO^ cells (Supplementary Table 1; Stroud et al., 2016). To confirm loss of the proteins of interest, mitochondria were isolated from control and two independent KO cell lines and analysed by western blotting (**Fig. 2A**). As expected, each KO cell line lacked expression of the targeted gene product. A dynamic interplay between the stability of other MCIA assembly factors was also observed. For example, in the absence of NDUFAF1, the levels of ECSIT and TMEM126B were strongly reduced relative to the control, while ACAD9 was unchanged (**Fig. 2A, lanes 1-3**). The ECSIT^KO^ cell lines mirrored this effect pointing to the close interaction between NDUFAF1 and ECSIT (**Fig. 2A, lanes 4- 6**). In the ACAD9^KO^ cells, the levels of NDUFAF1, ECSIT and TMEM126B were all reduced relative to the control cells (**Fig. 2A, lanes 7-9**). However, mitochondria lacking the membrane-integrated TMEM126B retained normal levels of other matrix facing MCIA complex proteins (**Fig. 2A, lanes 10- 15**). Taken together, these data suggest the stability of ACAD9 is independent of other known components of the MCIA complex, while the stability of TMEM126B is highly dependent on the presence of the NDUFAF1, ECSIT and ACAD9 (**Fig. 2B**). Loss of TIMMDC1 did not lead to changes in the steady-state levels of any of the MCIA complex subunits consistent with its role in the assembly of an independent complex I assembly module.

**Figure 2:**
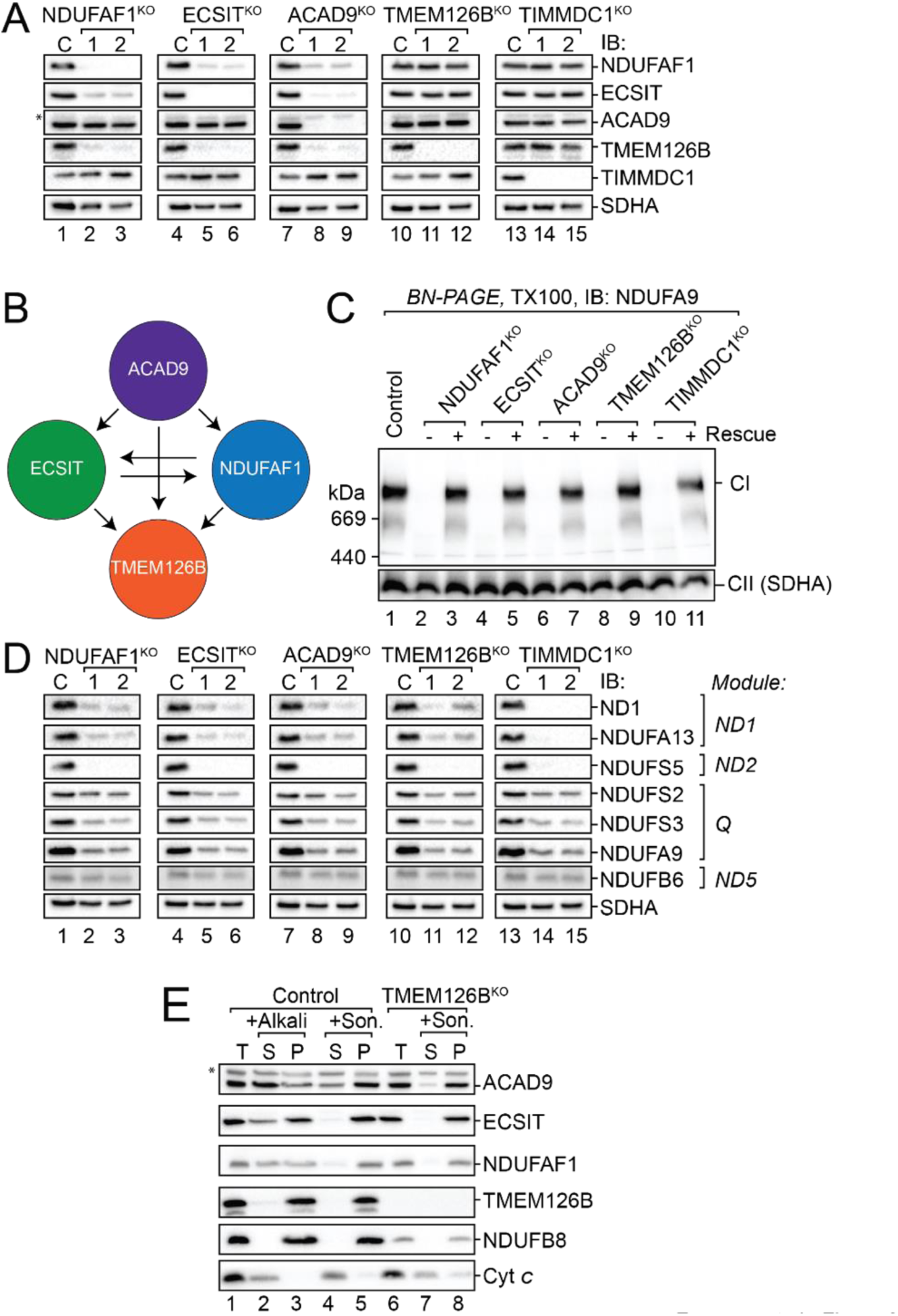
MCIA complex subunit hierarchy. **(A)** Mitochondrial proteins from control (C) and KO cell lines were subjected to SDS-PAGE and western blot analysis using antibodies as indicated. SDHA served as a loading control. **(B)** Schematic representation depicting the hierarchy of stability between MCIA complex components. **(C)** BN-PAGE and immunoblot analysis for complex I (NDUFA9). **(D)** SDS-PAGE and western blot analysis of complex I subunit levels in mitochondria from KO cells. **(E)** Mitochondria from control and TMEM126B^KO^ cells were subjected to alkali extraction or sonication with supernatant (S) and pellet (P) fractions analysed by SDS-PAGE and immunoblotting.

To assess how the loss of these assembly factors affect the steady-state levels of complex I, isolated mitochondria were solubilized in Triton X-100 and analysed by BN-PAGE and immunoblotting using antibodies against the complex I subunit NDUFA9. Loss of any individual assembly factor led to the complete loss of mature complex I (**Fig. 2C**). Furthermore, specificity of the genome editing was confirmed by complementation with the wild-type protein encoding a C-terminal Flag epitope (**Supplementary Fig. 1A**), with a complete rescue of the complex I defect observed in each case (**Fig. 2C**). Analysis of respiratory complexes III-V by BN-PAGE also showed normal levels relative to control mitochondria, with the only defect observed being the absence of the respirasome due to loss of complex I (**Supplementary Fig. 1B-F**). Next, we analysed the steady-state levels of representative complex I subunits in the absence of each assembly factor using SDS-PAGE and immunoblotting (**Fig. 2D**). Overall, there was a consistent reduction in the levels of complex I subunits relative to control mitochondria, with the intermembrane-space-localized NDUFS5 being the most strongly reduced subunit across all cell lines. Strong decreases were also observed for ND1, NDUFA13, NDUFS3, and NDUFA9 whereas NDUFS2 and NDUFB6 were moderately decreased. Residual amounts of NDUFA13 and mtDNA-encoded ND1 were detected in the absence of NDUFAF1, ECSIT, ACAD9 and TMEM126B, but no signal was observed in the TIMMDC1^KO^ cell lines (**Fig. 2D, compare ND1 and NDUFA13 lanes 1-12 to 13-15**). These data support TIMMDC1 having a prominent role in the biogenesis of ND1 and/or associated proteins, while the MCIA complex functions at a different stage of complex I assembly (Andrews et al., 2013; Guerrero-Castillo et al., 2017).

The interdependency of NDUFAF1, ECSIT, ACAD9 and TMEM126B led us to question if the multi-membrane spanning TMEM126B acts as the membrane anchor for the peripheral MCIA complex subunits as previously proposed (Guarani et al., 2014). In order to see if the remaining components of the MCIA complex were associated with the mitochondrial inner membrane in the absence of TMEM126B, a sonication and alkaline extraction was carried out on control and TMEM126B^KO^ mitochondria (**Fig. 2E**). In control cells, alkaline extraction could liberate a portion of ACAD9, ECSIT, NDUFAF1 and cytochrome *c* into the supernatant fraction following ultracentrifugation (**Fig. 2E, lane 2**), while the integral membrane proteins TMEM126B and NDUFB8 were present exclusively in the pellet fraction as expected (**Fig. 2E, lane 3**). Following sonication treatment of control mitochondria, cytochrome *c* was extracted into the soluble fraction (**Fig. 2E, lane 4**) while the remaining proteins were predominantly present in the pellet fraction (**Fig. 2E, lane 5**), demonstrating their association with the inner membrane. The same analysis performed on TMEM126B^KO^ mitochondria revealed that ACAD9, ECSIT and NDUFAF1 remained in the pellet fraction, similar to the membrane-embedded NDUFB8, while cytochrome *c* was extracted to the supernatant (**Fig. 2E, lanes 7 and 8**). These data indicate that in the absence of TMEM126B, the remaining components of the MCIA complex most likely remain membrane associated and that TMEM126B is not the exclusive membrane anchor of the MCIA complex.

### The MCIA steady-state complex is dependent on the presence of core ND2-module subunits

In order to gain insights into the behaviour of the MCIA complex upon loss of the individual components, we investigated steady-state ACAD9-containing complexes, since ACAD9 levels remained largely unchanged except in ACAD9^KO^ cells. In contrast to control mitochondria, ACAD9 complexes were absent in mitochondria lacking NDUFAF1 or ECSIT, with ACAD9 instead migrating as a low molecular weight species (**Fig. 3A**). In the TMEM126B^KO^ and TIMMDC1^KO^ cell lines, ACAD9 still formed high molecular weight complexes, however these appeared to differ in abundance and distribution relative to control mitochondria. We suggest that these represent crippled complex I intermediates containing ACAD9. In contrast, analysis of TIMMDC1 complexes revealed the stable accumulation of the 400/440 kDa intermediates along with the presence of a higher molecular weight complex in all MCIA KO mitochondria (**Fig. 3B**). Taken together, these results indicate that while all knockouts ultimately result in a complex I defect, NDUFAF1, ECSIT and ACAD9 are critical to forming all MCIA-dependent assembly intermediates, while loss of TMEM126B results in an altered assembly profile indicative of a perturbed assembly pathway.

**Figure 3:**
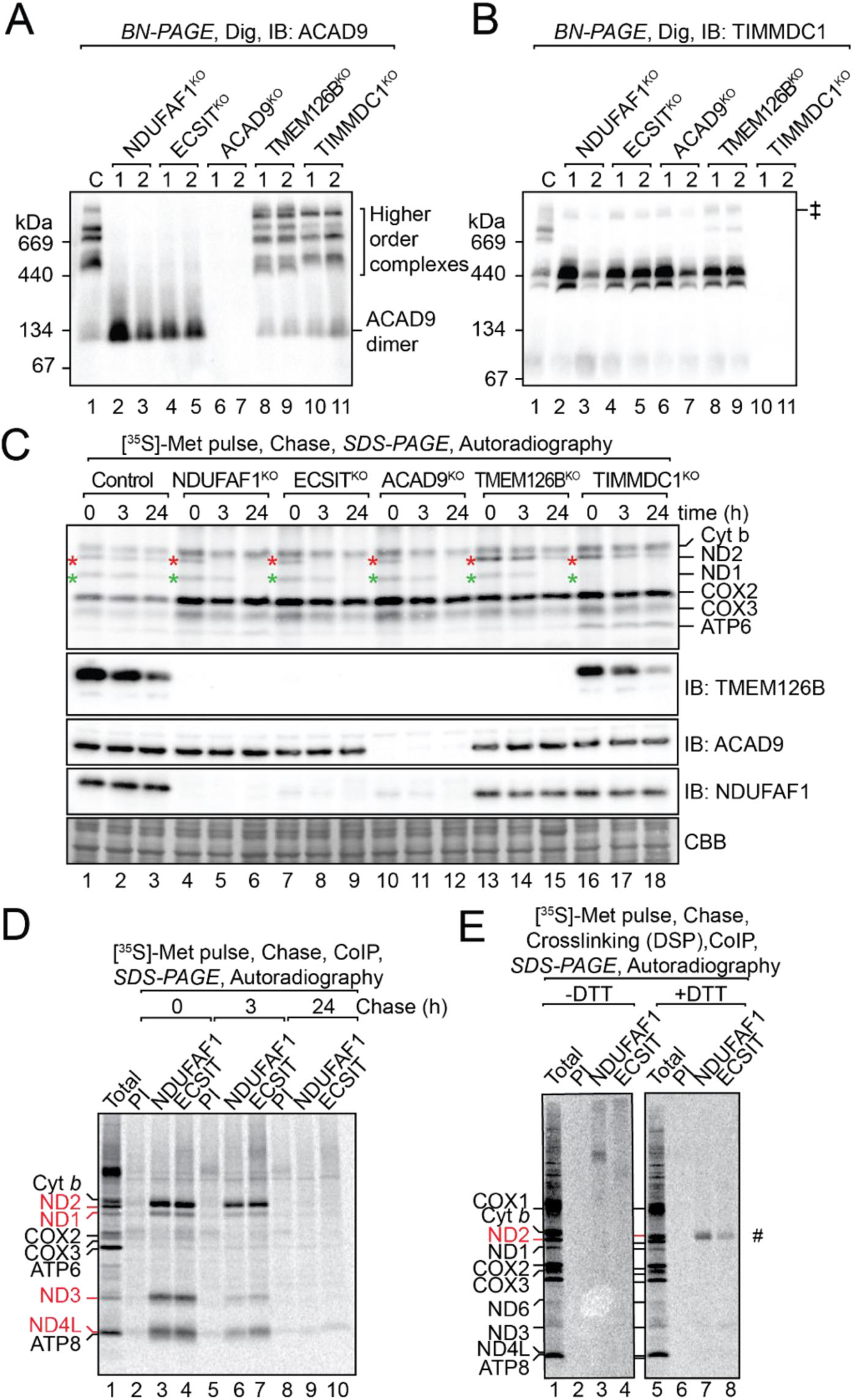
Loss of the MCIA complex alters complex I assembly intermediates. Mitochondria from indicated cell lines were solubilized in digitonin and analysed by BN-PAGE and immunoblotting (IB) for **(A)** ACAD9 and **(B)** TIMMDC1. **(C)** Cells were pulsed with [^35^S]-Met for 2 hours and chased for the indicated times. Isolated mitochondria were analysed by SDS-PAGE and autoradiography. ND1, green asterisk; ND2, red asterisk. IB for TMEM126B, ACAD9 and NDUFAF1 served as controls. Coomassie Brilliant Blue (CBB) staining was indicative of loading. **(D)** Following metabolic pulse chase labelling of 143B TK-cells, mitochondria were isolated and subjected to co-IP using antibodies as indicated. Elutions were then subjected to SDS-PAGE and autoradiography. **(E)** Samples (0h chase) as described in (D) were subjected to chemical crosslinking using DSP, followed by co-IP under denaturing conditions and analysis by SDS-PAGE under both reducing and non-reducing conditions and autoradiography. #, ND2.

Given the differences observed in subunit and assembly factor distribution in these KO cells, we sought to determine if there was a defect in the synthesis or stability of mtDNA-encoded subunits following pulse-chase metabolic labelling with [^35^S]-methionine (Formosa et al., 2016). Following the pulse-chase period, mitochondria were isolated and analysed by SDS-PAGE and autoradiography (**Fig. 3C**). In contrast to control cells, we observed turnover of ND2 in NDUFAF1^KO^, ECSIT^KO^ and ACAD9 ^KO^ cell lines by 3 hours post-chase and no signal detected by 24 hours (**Fig. 3C, lanes 3-12**). In TMEM126B^KO^ cells, newly synthesised ND2 was present at all time points, suggesting that TMEM126B is not required for ND2 stability over the 24-hour period (**Fig. 3C, lanes 13-15**). In NDUFAF1^KO^, ECSIT ^KO^, ACAD9 ^KO^ and TMEM126B^KO^ cells, newly translated ND1 levels were similar to that observed in control cells at the 0-hour timepoint (**Fig. 3C, compare lanes 1 with lanes 4**,**7**,**10 and 13**), however it was more readily degraded at the 3 and 24 hour timepoints (**Fig. 3C, lanes 4-15**). This was consistent with previous findings that loss of ND2 promoted increased turnover of ND1 (Dunning et al., 2007). In the TIMMDC1^KO^ cell line, ND2 was present and decreased in a similar manner to the control cells **(Fig. 3C, lanes 1-3 compared to lanes 16-18**), however the signal for ND1 was not detectable even directly following the pulse timepoint. This is consistent with findings that TIMMDC1 has a primary function in the translation or stability of ND1, while the remaining components of the MCIA complex is involved in the biogenesis of the ND2 module.

Since the ND2 module harbors mtDNA-encoded subunits ND2, ND3, ND4L and ND6, we investigated the interaction of NDUFAF1 and ECSIT with newly synthesised mtDNA-encoded subunits in further detail. Following pulse-chase labelling, mitochondria were isolated and subjected to co-immunoprecipitation using antibodies directed to the endogenous NDUFAF1 or ECSIT or pre-immune (PI) sera as a negative control **(Fig. 3D**). Directly after the pulse period (0h chase) there was a strong interaction of both NDUFAF1 and ECSIT with ND2 and to a lesser extent with ND1, ND3 and ND4L, which decreased at 3 hours post chase (**Fig. 3D; compare lanes 3 and 4 with lanes 6 and 7**). After 24 hours, where subunits have assembled into complex I, neither NDUFAF1 or ECSIT were found in association with mtDNA-encoded subunits (**Fig. 3D, lanes 9 and 10**). Importantly, the pre-immune sera did not enrich any subunits (**Fig. 3D; lanes 2, 5 and 8**). Given the relationship of the MCIA complex with newly synthesised ND2, we sought to further investigate how mitochondrial translation inhibition affects the stability and assembly of this assembly complex. We analysed 143B control (ρ^+^) and ρ^0^ cells, which lack mtDNA, as well as control HEK293T cells grown in the presence of chloramphenicol (CAP) to inhibit mitoribosome translation. Analysis of these two treatments showed indistinguishable phenotypes when analysed by SDS-PAGE and BN-PAGE (**Supplementary Fig. 2**). We observed that NDUFAF1, ECSIT and TMEM126B protein levels were reduced relative to their respective controls, as were the complex I subunits NDUFA13 and NDUFC2 while ACAD9 was largely unaffected **(Supplementary Fig. 2A**). Consistent with this, reduction of MCIA complexes was also observed (**Supplementary Fig. 2B and 2D**). These data suggest that the stability of the MCIA complex and its constituents is also dependent on the translation of associated mtDNA-encoded subunits. To differentiate between direct protein-protein interactions or associative interactions within a complex, chemical crosslinking using a thiol-cleavable crosslinker dithiobis-succinimidyl propionate (DSP) and co-immunoprecipitation was performed on mitochondrial lysates under denaturing conditions (**Fig. 3E**). Following crosslinking under oxidizing conditions (-DTT), a clear radiolabelled species was observed in the NDUFAF1 elution samples that was absent in the pre-immune control (**Fig. 3E; lanes 2 and 3**). Under reducing conditions where the crosslinker is cleaved (+DTT), the radiolabelled species observed following elution from NDUFAF1 antibodies was revealed to be ND2, which was also present to a lesser extent after elution from ECSIT antibodies (**Fig. 3E, #**). Taken together, these data suggest that newly-synthesised ND2 may be translated independently of the MCIA complex before making direct contacts with NDUFAF1 and ECSIT for further assembly. In the absence of these assembly factors the protein is unstable and degraded.

### Identification of TMEM186 as a component of the MCIA complex

Through analysis of the MCIA components it became clear that the ∼450kDa complex observed on BN-PAGE could not be fully accounted for by the molecular mass of the known protein constituents. To address this, we utilized proximity-dependent biotin identification (BioID)(Roux et al., 2012) to label and isolate proteins in proximity to the MCIA complex. NDUFAF1^KO^ cells were rescued with NDUFAF1 fused to the promiscuous biotin ligase BirA* (NDUFAF1^BirA*^) and biotinylated proteins detected by quantitative mass spectrometry (Q-MS). All known components of the MCIA complex were found in proximity to the BirA*, as well as a number of complex I subunits enriched in the NDUFAF1^BirA*^ sample (**Fig. 4A; Supplementary Table 2**). TMEM186 was also found significantly enriched in the NDUFAF1^BirA*^ sample. Recently, TMEM186 was found to co-migrate with components of the MCIA complex using dynamic complexome profiling of BN-PAGE gel slices (Guerrero-Castillo et al., 2017). However, the importance of this protein in the assembly of complex I has not been addressed. *In vitro* import analysis revealed that TMEM186 contains a cleavable mitochondrial presequence and its import is dependent on the mitochondrial membrane potential (**Fig. 4B**). TMEM186 is predicated to have two transmembrane anchors (Krogh et al., 2001). Alkaline extraction studies of mitochondria from cells expressing TMEM186 with a C-terminal Flag epitope (TMEM186^Flag^) revealed that TMEM186^Flag^ behaves like integral membrane protein Mic10, in contrast to peripheral membrane proteins NDUFAF1 and NDUFS2 and cytochrome *c* (**Fig. 4C**). Furthermore, submitochondrial analysis using protease accessibility revealed that TMEM186^Flag^ was protected following outer membrane rupture, indicating that the C-terminal Flag faces the matrix (**Fig. 4D**). Mitochondrial matrix resident NDUFAF1 and NDUFS2 were also protected from external protease following swelling while the intermembrane space-accessible protein Mic10 was only degraded following outer membrane rupture. As expected, the cytosolic exposed Mfn2 was degraded both with and without swelling (**Fig. 4D lanes 2 and 4**). Taken together, these data suggest that TMEM186 is inserted into the mitochondrial inner membrane, with both the N- and C-termini facing the matrix (**Fig. 4E**).

**Figure 4:**
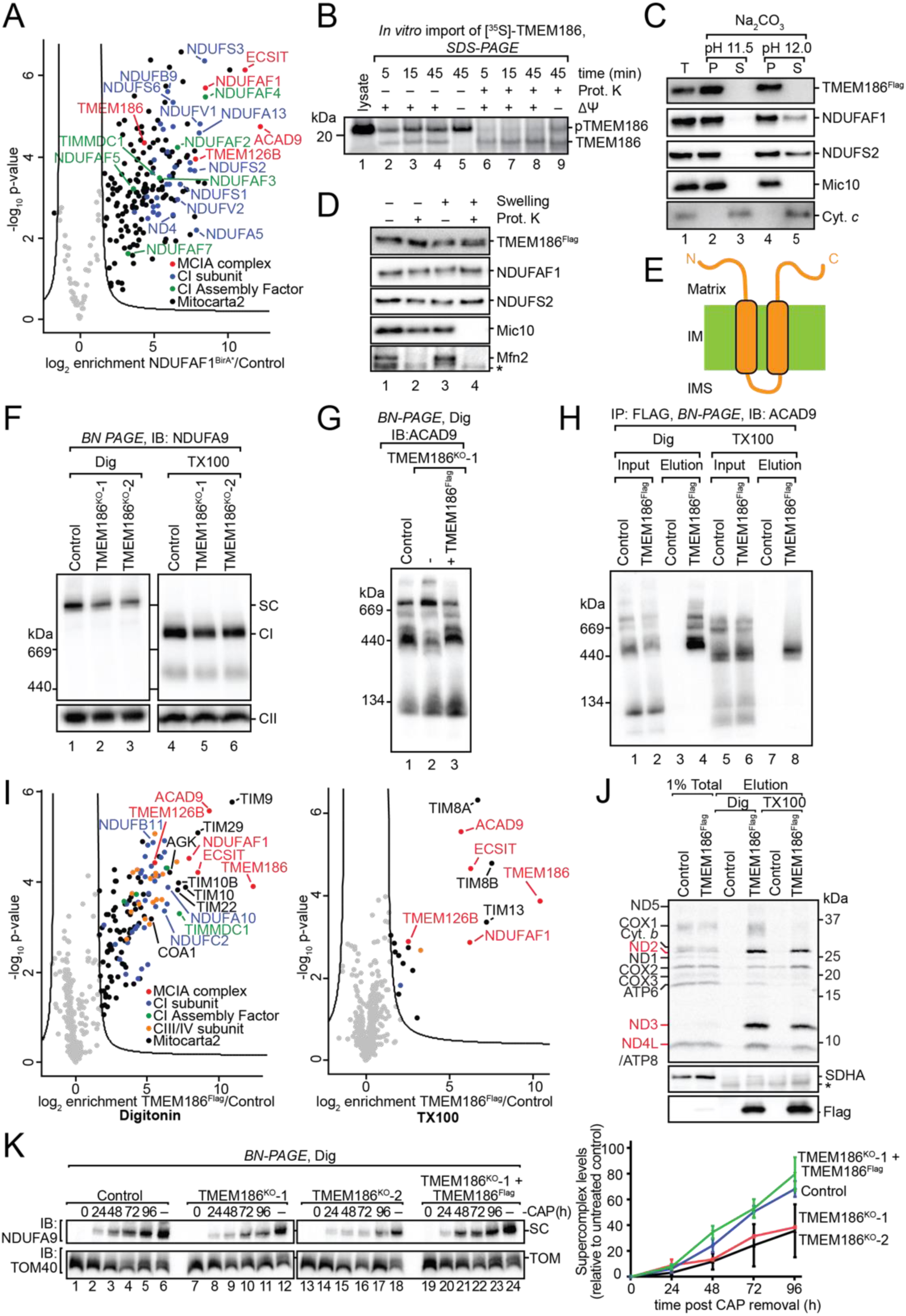
TMEM186 is a component of the MCIA complex. **(A)** BioID proximity labelling and protein enrichment in NDUFAF1^BirA*^ relative to control. **(B)** *In vitro* import analysis of [^35^S]-TMEM186. **(C)** Mitochondria from cells expressing TMEM186^Flag^ were subjected to alkali extraction (Na2CO3) followed by SDS-PAGE and western blot analysis. **(D)** Mitochondria from cells expressing TMEM186^Flag^ were subjected to a swelling and protease protection assay followed by proteinase K (Prot. K) incubation where indicated. **(E)** Schematic representation of TMEM186 topology. **(F)** Control and TMEM186^KO^ mitochondria were solubilized in digitonin or Triton X-100, followed by BN-PAGE and immunoblotting with NDUFA9 antibodies. Complex II (SDHA) served as a loading control. **(G)** Control, TMEM186^KO^ or rescue (TMEM186^KO^+TMEM186^Flag^) mitochondria were solubilized in digitonin followed by BN-PAGE and immunoblotting with ACAD9 antibodies. **(H)** Digitonin or Triton X-100 solubilized control or TMEM186^KO^ +TMEM186^Flag^ mitochondria were subjected to affinity enrichment using Flag agarose beads, followed by BN-PAGE analysis and **(I)** label-free quantitative proteomics. **(J)** Following mtDNA-encoded subunit labelling, mitochondria were isolated and subjected to Flag affinity enrichment. Eluates were analysed by SDS-PAGE and phosphorimaging. Immunoblotting with Flag and SDHA antibodies served as controls. **(K)** Following chloramphenicol (CAP) treatment, cells were chased and complex I *de novo* assembly analysed by BN-PAGE. TOM40 served as a loading control. Right panel: Quantitation of NDUFA9 signal normalized to TOM40, all relative to untreated control (%). Error bars: n=3, mean ± SEM. Asterisk (*) indicates a non-specific band.

We generated KO cell lines of TMEM186 (**Supplementary Table 1**). Analysis of complex I from TMEM186^KO^ cell lines by BN-PAGE showed only a modest decrease in levels – both in the supercomplex form (using digitonin solubilization) and holoenzyme form (using Triton X-100 solubilization) (**Fig. 4F**). Analysis of the steady state MCIA complexes revealed reduced levels, and faster migration of the lowest MCIA complex that was rescued upon re-expression of TMEM186^Flag^ (**Fig. 4G and Supplementary Fig. 3A, lanes 1-3**). In contrast, TIMMDC1 complexes were unaffected (**Supplementary Fig. 3A, lanes 4-6**). Complexes III, IV and V showed no striking differences in the steady-state levels between control and TMEM186^KO^ mitochondria (**Supplementary Fig. 3B**).

In order to determine if TMEM186 is a constituent of the MCIA complex, we solubilized control or TMEM186^Flag^ mitochondria in digitonin or Triton X-100 and performed immuno-enrichment of TMEM186^Flag^ complexes using Flag beads, followed by elution under native conditions and subsequent analysis by BN-PAGE (**Fig. 4H**). Using mild digitonin solubilization, TMEM186^Flag^ was able to enrich the higher molecular weight ACAD9-containing complexes but not unassembled ACAD9 (**Fig. 4H**). Under the more stringent, but still native conditions of Triton X-100 solubilization, TMEM186^Flag^ was only able to enrich the ∼450kDa complex, suggesting a stronger interaction with this assembly intermediate. The elutions were also subjected to Q-MS analysis for an unbiased overview of the protein interaction landscape. The digitonin solubilized TMEM186^Flag^ elutions revealed the significant enrichment of numerous proteins including a number of complex I subunits and assembly factors, as well as some subunits of respiratory complexes III and IV and the TIM22 translocase complex (**Fig. 4I, left panel; Supplementary Table 3**). The most highly enriched proteins were MCIA assembly factors NDUFAF1, ECSIT and ACAD9, and to a lesser extent TMEM126B. In Triton X-100 eluates, TMEM186 specifically enriched NDUFAF1, ECSIT and ACAD9 (**Fig. 4I, right panel; Supplementary Table 4**) whereas complex I, III and IV subunits were not strongly enriched. The enrichment of TMEM186 with the TIM22 machinery (TIM22, TIM29, AGK, small TIMs) may be due to TMEM186 using the TIM22 import pathway for membrane integration (Callegari et al., 2016; Kang et al., 2016; Kang et al., 2018; Kang et al., 2017; Vukotic et al., 2017) as well as overexpression of the Flag tagged protein.

To determine if TMEM186 can associate with newly synthesised mtDNA-encoded subunits, control and TMEM186^Flag^ expressing cells were pulse labelled with [^35^S]-Methionine and subjected to co-immunoprecipitation using Flag beads. As can be seen (**Fig. 4J**), ND3 was the most enriched subunit under both digitonin and Triton X-100 solubilization conditions. TMEM186^Flag^ also specifically enriched ND2 and ND4L. We conclude that TMEM186 indeed interacts with the core MCIA members NDUFAF1, ECSIT and ACAD9, as part of the biogenesis of the ND2 module with a possible role in biogenesis of the ND3 subunit.

Because the loss of TMEM186 does not block complex I biogenesis, we decided to investigate whether the rate of *de novo* complex I assembly was altered in the KO cells. Steady-state respiratory complexes were depleted from control, TMEM186^KO^ clones and rescue cell lines by CAP treatment for 96 hours. CAP was then removed and *de novo* assembly of complex I analysed (**Fig. 4K**). In both TMEM186^KO^ cell lines, the rate of complex I assembly was decreased to ∼40% of control mitochondria, while this defect was rescued upon expression of TMEM186^Flag^ (**Fig. 4K**). We conclude that while TMEM186 is not an essential component of the complex I assembly machinery, loss of TMEM186 is refractory to the optimal function of the MCIA complex resulting in reduced assembly of complex I.

### COA1 is required for complex I assembly, but dispensable for complex IV assembly

Guerrero-Castillo and colleagues (2017) recently proposed that the previously reported complex IV assembly factor COA1 (also known as MITRAC15 (Mick et al., 2012)) may play a role in the biogenesis of complex I since it co-migrated on BN-PAGE with components of the MCIA complex and assembly intermediates. Consistent with this, we found that under mild conditions of digitonin solubilization, COA1 was also enriched with TMEM186^Flag^ (**Fig. 4I**). As yet, the importance of COA1 in complex I assembly has yet to be specifically addressed. We generated and validated COA1^KO^ cells (**Fig. 5A; Supplementary Table 1**). BN-PAGE analysis revealed that complex I still assembled in the COA1^KO^ cell lines but at a ∼50% reduction in comparison to control mitochondria (**Fig. 5B**). In contrast to previous reports using COA1 depletion by RNA interference (Mick et al., 2012), knockout of COA1 did not appear to have an effect on complex IV levels (**Fig. 5C; Supplementary Fig. 4A**). No differences were observed in the levels of complex III **(Supplementary Fig. 4B**) or complex V (**Supplementary Fig. 4C**).

**Figure 5:**
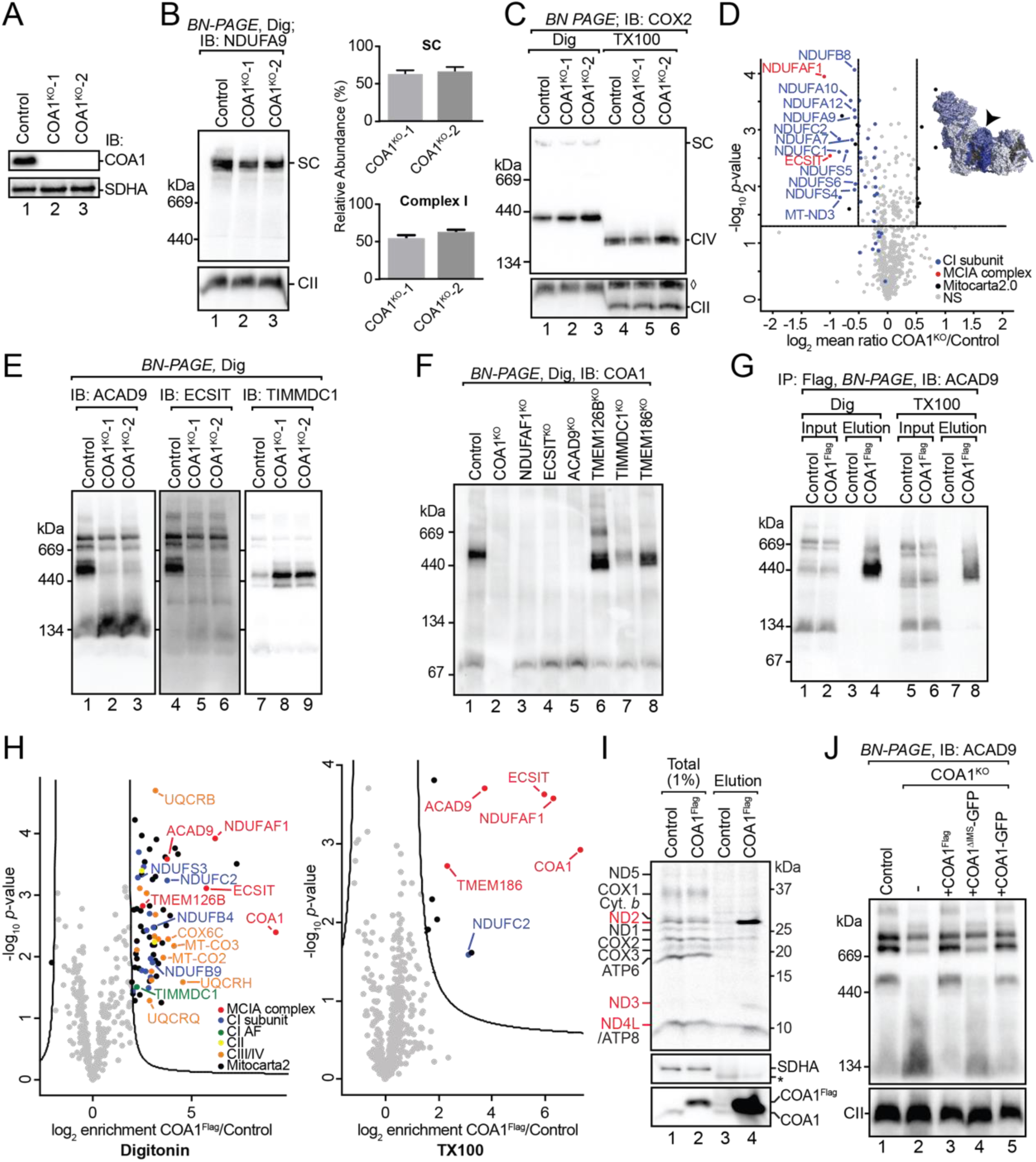
COA1 participates in complex I assembly and is required for stability of the MCIA complex. **(A)** SDS-PAGE and western blot analysis of COA1 in mitochondria from control and COA1^KO^ cells. SDHA served as a loading control. **(B)** Mitochondria from control and COA1^KO^ cells were solubilized in digitonin and subjected to BN-PAGE and western blotting using NDUFA9 antibodies. Complex II (CII, SDHA) served as a loading control. Relative abundance to control supercomplex (SC) or complex I shown as mean ± SEM, n=3. **(C)** Mitochondria from control and COA1^KO^ cells were solubilized in digitonin or Triton X-100 and subjected to BN-PAGE and western blotting using COX2 antibodies. Complex II (CII, SDHA) served as a loading control. ◊ indicates previous COX2 signal. **(D)** Volcano plot showing proteins with altered abundance in COA1^KO^ mitochondria. Inset: Complex I subunit changes mapped onto the structure of complex I. **(E)** Mitochondria from control and COA1^KO^ cells were solubilized in digitonin and subjected to BN-PAGE and western blotting using antibodies directed to ACAD9, ECSIT and TIMMDC1. **(F)** Mitochondria from control and MCIA KO cells were solubilized in digitonin and subjected to BN-PAGE and western blotting using COA1 antibodies. **(G)** Control or COA1^KO^+COA1^Flag^ mitochondria were subjected to detergent solubilization with digitonin or Triton X-100 and affinity enrichment using Flag agarose beads. Elutions were subjected to BN-PAGE and western blotting using ACAD9 antibodies. **(H)** Samples were prepared as in (G) and subjected to LFQ proteomics. **(I)** Pulse labelled mtDNA-encoded subunits from control or COA1^KO^+COA1^Flag^ cells were subjected to Flag affinity enrichment. Eluates were analysed by SDS-PAGE and phosphorimaging. Immunoblotting with COA1 and SDHA antibodies served as controls. **(J)** Mitochondria from control, COA1^KO^ or COA1^KO^ cells expressing COA1^Flag^, COA1-GFP^Flag^ and COA1^ΔIMS^-GFP^Flag^ were solubilized in digitonin and subjected to BN-PAGE and immunoblotting for ACAD9.

Given the specific decrease in complex I levels, we performed SILAC and quantitative mass spectrometric (Q-MS) analysis and found reduced levels of NDUFAF1 and ECSIT in COA1^KO^ mitochondria as well as complex I subunits belonging to the ND2-module (**Fig. 5D; Supplementary Table 5**), consistent with defects in the MCIA complex. This was recapitulated using SDS-PAGE and immunoblotting of mitochondria from control and COA1^KO^ cells for various proteins (**Supplementary Fig. 4D**), showing decreased protein levels for TMEM126B (not quantified by Q-MS), ECSIT and NDUFAF1, and complex I subunits NDUFC2 and NDUFS5. No changes to ACAD9, TIMMDC1, the complex I subunit NDUFS2, or complex IV subunits COX1 and COX4 were observed. We next addressed whether loss of COA1 affects the assembly or stability of the MCIA complex (**Fig. 5E**). The ∼450kDa MCIA complex was specifically lost with the accumulation of free ACAD9 but not ECSIT (**Fig. 5E**). Similar to what was observed for other MCIA knockout cell lines, loss of COA1 also resulted in the accumulation of TIMMDC1 complexes containing the ND1/Q modules (**Fig. 5E**).

To further investigate how COA1 behaves with respect to the MCIA complex, we performed BN-PAGE using mitochondria from each of the MCIA complex KO lines (**Fig. 5F**). In control cells, COA1 assembled in a ∼450kDa complex, which was absent in cells lacking NDUFAF1, ECSIT or ACAD9, thus pointing to a direct and stable association of COA1 with this complex. Supporting this, COA1 migrated in the MCIA complexes that were altered in size or abundance in TMEM126B^KO^, TIMMDC1^KO^ and TMEM186^KO^ mitochondria (**Fig. 5F**). We conclude that COA1 is integral to the stabilisation of the ∼450kDa MCIA complex.

Expression of COA1^Flag^ in COA1^KO^ cells restored complex I (**Supplementary Fig. 4E**), ACAD9 and TIMMDC1 complexes back to control levels (**Supplementary Fig. 4F**). Co-immunoprecipitation analysis under both digitonin and Triton X-100 solubilization conditions, resulted in the enrichment of the ∼450kDa MCIA species (**Fig. 5G**). Analysis of COA1^Flag^ elutions by Q-MS revealed that under mild digitonin conditions, COA1 highly enriched NDUFAF1, ECSIT and ACAD9 along with a number of respiratory chain subunits from complexes I, III and IV (**Fig. 5H, left panel; Supplementary Table 6**). Q-MS analysis of Triton X-100 solubilized mitochondria supported a stable interaction between COA1 and MCIA components ACAD9, ECSIT, NDUFAF1 and TMEM186 (**Fig. 5H, right panel; Supplementary Table 7**). In addition, NDUFC2, a complex I subunit belonging to the ND2 module (Fiedorczuk et al., 2016; Stroud et al., 2016) was also enriched. To investigate the possible interaction between COA1 and newly translated mtDNA-encoded subunits, we performed co-immunoprecipitation analysis of pulse-labelled mtDNA translation products. In this case, COA1 efficiently immunoprecipitated newly synthesised ND2, and to a lesser extent ND3 and ND4L (**Fig. 5I**). Taken together, COA1 is a structural component of the MCIA complex and is critical to the formation and stability of the ∼450kDa assembly intermediate complex and the ND2 module.

In comparison to other components of the MCIA complex, COA1 has an unusual topology, with only a small matrix facing polypeptide (∼14 aa) followed by a single transmembrane anchor and a ‘Tim21-family’ domain facing the IMS (Mick et al., 2012). To determine if the IMS domain is required for COA1 function, we generated COA1^KO^ cell lines expressing either COA1 with a C-terminal Flag tag (COA1^Flag^), as a GFP fusion (COA1-GFP), or with the transmembrane anchor of COA1 (amino acids 1-37) attached to GFP and hence lacking the IMS domain (COA1^ΔIMS^-GFP) (**Supplementary Fig. 4G**). Cells expressing COA1-GFP could restore assembly of ACAD9 complexes to the same degree as COA1^Flag^, but loss of the IMS domain did not (**Fig. 5J**). We conclude that while topologically separated from the remaining components of the MCIA complex, the IMS domain of COA1 is important for the biogenesis of complex I through the stabilisation of the ∼450 kDa MCIA complex. These data suggest a prominent role of COA1 in the assembly of complex I through the MCIA complex.

## Discussion

### Dissection of the MCIA complex components

The precise role of many complex I assembly factors is poorly understood, with suggested roles in subunit maturation, assembly and stabilisation of specific modules and in the integration of modules into higher ordered assemblies (Formosa et al., 2018). It appears that most complex I assembly factors fall under the umbrella of chaperones that stabilize assembly intermediates. However, two have been shown to perform post-translational modification (Rhein et al., 2013; Rhein et al., 2016) and another in the delivery of cofactors to specific subunits (Sheftel et al., 2009).

Our analysis of complex I subunit knockout lines are consistent with previous studies (Guerrero-Castillo et al., 2017; Pagliarini et al., 2008; Ugalde et al., 2004) that describe distinct complex I intermediate assemblies that accumulate and further associate into larger subcomplexes. Loss of functional subassemblies also leads to the formation of non-productive, crippled intermediates (Formosa et al., 2015). The MCIA complex represents a key factor in the assembly of complex I and its importance has long been appreciated with analysis of both patient derived fibroblasts as well as knockdown in cell culture (Dunning et al., 2007; Heide et al., 2012; Nouws et al., 2010; Vogel et al., 2005; Vogel et al., 2007b). Furthermore, homologues of some MCIA complex assembly factors exist in most organisms that harbour complex I, including *Arabidopsis thaliana, Neurospora crassa, Caenorhabditis elegans* and *Drosophila melanogaster* (Chuaijit et al., 2019; Garcia et al., 2017; Kuffner et al., 1998; Ligas et al., 2019). What has remained unclear is the interplay of the components of the MCIA complex and how they each may contribute to assembly.

Here, we found that deletion of each component individually leads to (1) different effects on MCIA complex subunit levels; (2) different steady-state levels of intermediate complexes; and (3) differences in the assembly kinetics of complex I. ACAD9 is required for the stability of all other components of the MCIA complex, while TMEM126B is not a requisite for stability of the remaining subunits of the assembly complex. Furthermore, while ACAD9 protein levels were unchanged upon loss of NDUFAF1 and ECSIT, both of these assembly factors are essential for ACAD9 to form higher molecular weight complexes. Taking into consideration the interconnected stability and effect on the assembly pathway, ACAD9-ECSIT-NDUFAF1 appear to form the ‘core’ of the MCIA complex, while TMEM126B may engage more transiently. A number of critical observations lead to this conclusion. Firstly, loss of TMEM126B does not preclude the formation of higher-order molecular weight complexes containing ACAD9, which are critically dependent on NDUFAF1 and ECSIT. It has also been previously demonstrated that deletion of ECSIT in bone marrow-derived macrophages similarly result in decreased NDUFAF1 levels (Carneiro et al., 2018). Secondly, TMEM126B was not readily isolated with COA1 or TMEM186 under the harsher condition imposed by Triton X-100 solubilization, whereas NDUFAF1, ECSIT and ACAD9 were consistently enriched. Finally, deletion of TMEM126B did not affect the stability of the remaining MCIA complex components, suggesting that they are capable of forming a stable complex. It should be noted that this in no way reduces the importance of TMEM126B in the assembly process as loss of the protein blocks complex I assembly. Indeed, modulation of TMEM126B levels may also be a mechanism for direct regulation of complex I levels, for example during chronic hypoxia (Fuhrmann et al., 2018). In this case, the other MCIA components would still be present but may be non-functional.

### Assembly and stability of the MCIA complex is coupled to translation

The MCIA complex is dependent on the expression of mtDNA. TMEM126B, ECSIT and NDUFAF1 levels were strongly reduced upon depletion of mtDNA or by inhibition of mitochondrial translation. The levels of ACAD9 were largely unaffected, consistent with its stoichiometric excess and proposed moonlighting role in beta-oxidation of fatty acids (Nouws et al., 2013). From our results, we suggest that the assembly of the MCIA complex is stabilised by the presence of newly translated subunits – in particular that of ND2. This is supported by the direct interaction of ND2 with NDUFAF1 and ECSIT through chemical cross-linking as well as previous complexome studies into de novo complex I assembly (Guerrero-Castillo et al., 2017). Previous quantitative proteomic analysis of cells lacking the ND2 module accessory subunits NDUFC1 or NDUFC2 subunits also did not show decreases in the levels of MCIA subunit proteins (Stroud et al., 2016) consistent with the observation of higher molecular weight complexes containing ACAD9 in these cells.

### TMEM186 and COA1 are MCIA complex subunits required for efficient assembly of complex I

We also demonstrated that TMEM186 and COA1 are components of the MCIA complex where they appear to play more peripheral functions to the core machinery. Until now, little was known regarding the role of TMEM186 other than an interaction with ECSIT (Guarani et al., 2014) and co-migration with the MCIA complex on BN-PAGE (Guerrero-Castillo et al., 2017). Here we show that deletion of TMEM186 does not appear to affect the steady-state levels of complex I, but rather the rate of *de novo* assembly of the enzyme. This is the first indication that an assembly factor may influence the rate of assembly, which adds to the repertoire of functions these proteins perform. We also observed a strong enrichment of newly-translated ND3 associated with TMEM186. Such enrichment of ND3 was observed for other subunits of the MCIA complex, however not to the same extent. This is in agreement with the assembly model build based on complexome profiling suggesting that TMEM186 and ND3 enter the assembly pathway at the same stage (Guerrero-Castillo et al., 2017). The reduced kinetics of complex I assembly in TMEM186^KO^ cells may be due to perturbed ND3 integration at this step.

Previous studies indicated that COA1 is a component of the MITRAC complex involved in the assembly or stability of the complex IV subunit COX1, however a reduction in complex I levels was also shown upon COA1 knockdown (Mick et al., 2012). A possible role in complex I assembly was more recently suggested since COA1 co-migrated with the MCIA complex and complex I intermediates on BN-PAGE (Guerrero-Castillo et al., 2017). Through knockout approaches, we were able to show that neither the level of assembled complex IV nor its constituent subunits appeared to be affected by deletion of COA1. This raises the question what COA1 may be doing in association with complex IV. It also cannot be ruled out that deletion of COA1 may present differently in different tissues and a complex IV defect may be observed in a tissue-dependent context. Another interesting aspect of COA1 function is the peculiar topology this protein exhibits, where it contains an N-terminal transmembrane anchor and a domain that protrudes into the intermembrane space (Mick et al., 2012). We found that the IMS domain is critical for the stability of the MCIA complex. So how does this domain, which is spatially segregated from the bulk of the MCIA complex, help in co-ordinating complex I assembly? The answer may lie in the strong interaction observed between newly synthesized ND2 and COA1. ND2 contains 10 transmembrane-spanning helices and needs to be carefully threaded into the membrane. Newly synthesised ND2 first accumulates in the ∼450kDa complex (Dunning et al., 2007; Lazarou et al., 2007) where it also cross links with ECSIT and NDUFAF1. This complex was also strongly destabilized in the COA1^KO^ cells. Future work into how COA1 functions mechanistically will be crucial to untangle the role of this protein.

How defects in TMEM186 and COA1 would present in a physiological or disease context presents an intriguing question. As no patients have yet been reported with mitochondrial dysfunction resulting from mutations in either of these genes, one would predict that complex I deficiency may present in tissues where complex I assembly is slow or where mitochondrial turnover is high. Nevertheless, the identification of TMEM186 and COA1 as *bona fide* assembly factors for complex I presents a new avenue for the identification of patients that present with complex I deficiency.

## Author Contributions

Conceptualization, L.E.F. and M.T.R; Methodology, L.E.F., L.M.W., B.R., A.J.S., M.L., D.A.S. and M.T.R; Formal Analysis, L.E.F., B.R., D.A.S and M.T.R.; Investigation, L.E.F., L.M.W., B.R., A.J.S., and M.L.; Data Curation, L.E.F., B.R, D.A.S. and M.T.R; Writing – Original Draft, L.E.F. and M.T.R; Writing – Review & Editing, L.E.F., L.M.W., B.R., A.J.S., T.H.B., M.L., D.A.S. and M.T.R; Visualization, L.E.F. and M.T.R; Supervision, D.A.S. and M.T.R.; Project Administration, M.T.R; Funding Acquisition, D.A.S. and M.T.R.

## Acknowledgments

We thank Dr Felix Kraus and Dr Marris Dibley for constructive discussions. We thank the Bio21 Mass Spectrometry and Proteomics Facility (MMSPF) and the Monash Biomedical Proteomics Facility for the provision of instrumentation, training, and technical support and Monash Flowcore for cell sorting. We acknowledge funding from the National Health and Medical Research Council (NHMRC Project Grants 1164459 to MTR; 1125390, 1140906 to MTR and DAS; NHMRC Fellowship 1140851 to DAS).

## Methods and materials

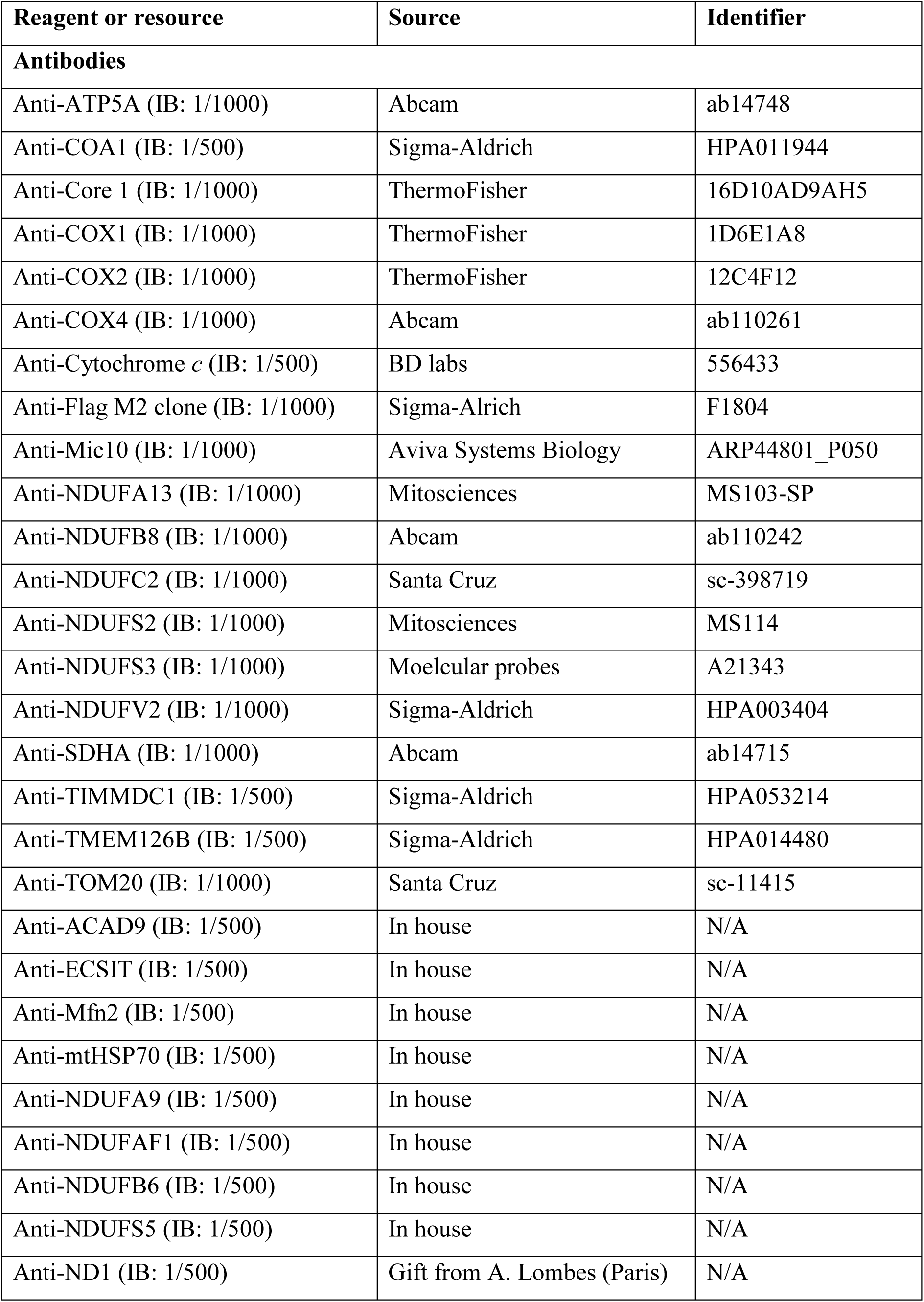
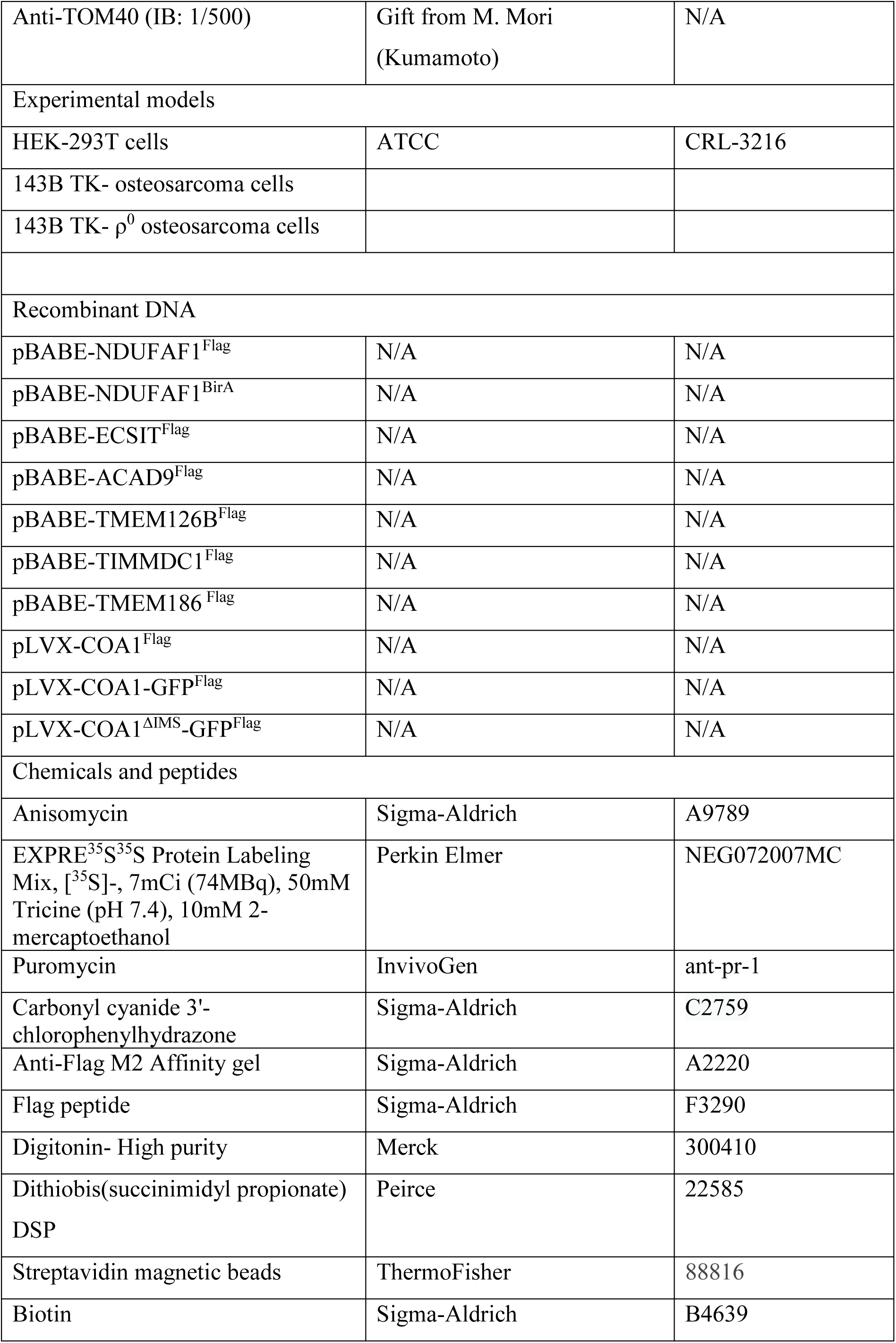
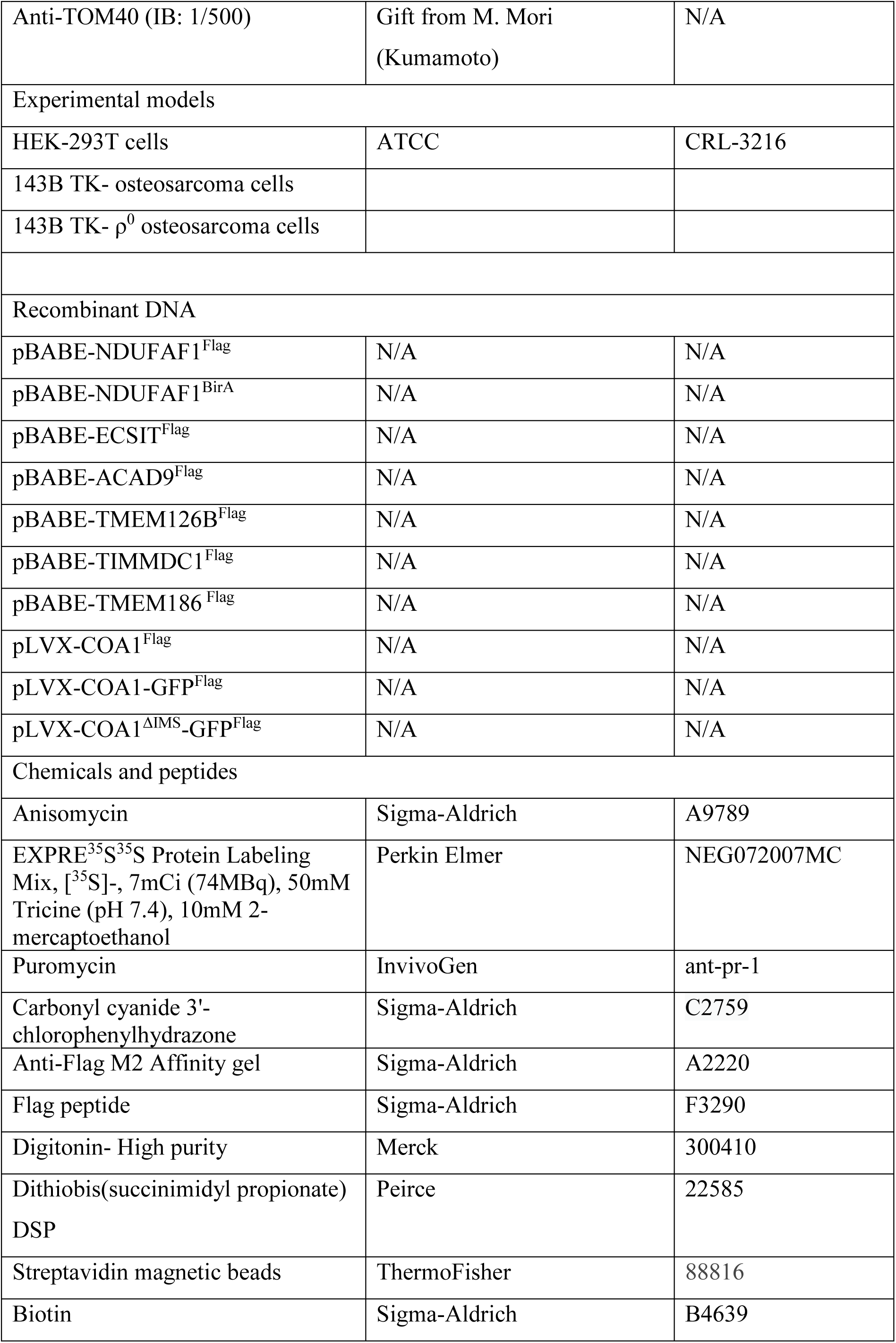
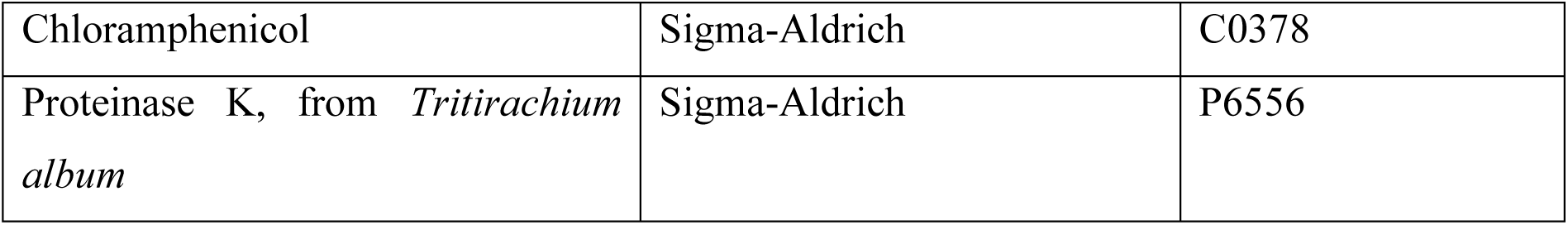

| Reagent or resource | Source | Identifier |
| --- | --- | --- |
| **Antibodies** |  |  |
| Anti-ATP5A (IB: 1/1000) | Abcam | ab14748 |
| Anti-COA1 (IB: 1/500) | Sigma-Aldrich | HPA011944 |
| Anti-Core 1 (IB: 1/1000) | ThermoFisher | 16D10AD9AH5 |
| Anti-COX1 (IB: 1/1000) | ThermoFisher | 1D6E1A8 |
| Anti-COX2 (IB: 1/1000) | ThermoFisher | 12C4F12 |
| Anti-COX4 (IB: 1/1000) | Abcam | ab110261 |
| Anti-Cytochrome *c* (IB: 1/500) | BD labs | 556433 |
| Anti-Flag M2 clone (IB: 1/1000) | Sigma-Alrich | F1804 |
| Anti-Mic10 (IB: 1/1000) | Aviva Systems Biology | ARP44801_P050 |
| Anti-NDUFA13 (IB: 1/1000) | Mitosciences | MS103-SP |
| Anti-NDUFB8 (IB: 1/1000) | Abcam | ab110242 |
| Anti-NDUFC2 (IB: 1/1000) | Santa Cruz | sc-398719 |
| Anti-NDUFS2 (IB: 1/1000) | Mitosciences | MS114 |
| Anti-NDUFS3 (IB: 1/1000) | Moelcular probes | A21343 |
| Anti-NDUFV2 (IB: 1/1000) | Sigma-Aldrich | HPA003404 |
| Anti-SDHA (IB: 1/1000) | Abcam | ab14715 |
| Anti-TIMMDC1 (IB: 1/500) | Sigma-Aldrich | HPA053214 |
| Anti-TMEM126B (IB: 1/500) | Sigma-Aldrich | HPA014480 |
| Anti-TOM20 (IB: 1/1000) | Santa Cruz | sc-11415 |
| Anti-ACAD9 (IB: 1/500) | In house | N/A |
| Anti-ECSIT (IB: 1/500) | In house | N/A |
| Anti-Mfn2 (IB: 1/500) | In house | N/A |
| Anti-mtHSP70 (IB: 1/500) | In house | N/A |
| Anti-NDUFA9 (IB: 1/500) | In house | N/A |
| Anti-NDUFAF1 (IB: 1/500) | In house | N/A |
| Anti-NDUFB6 (IB: 1/500) | In house | N/A |
| Anti-NDUFS5 (IB: 1/500) | In house | N/A |
| Anti-ND1 (IB: 1/500) | Gift from A. Lombes (Paris) | N/A |
| Anti-TOM40 (IB: 1/500) | Gift from M. Mori<br>(Kumamoto) | N/A |
| Experimental models |  |  |
| HEK-293T cells | ATCC | CRL-3216 |
| 143B TK- osteosarcoma cells |  |  |
| 143B TK- $\rho^0$ osteosarcoma cells | | |
| Recombinant DNA |  |  |
| pBABE-NDUFAF1<sup>Flag</sup> | N/A | N/A |
| pBABE-NDUFAF1<sup>BirA</sup> | N/A | N/A |
| pBABE-ECSIT<sup>Flag</sup> | N/A | N/A |
| pBABE-ACAD9<sup>Flag</sup> | N/A | N/A |
| pBABE-TMEM126B<sup>Flag</sup> | N/A | N/A |
| pBABE-TIMMDC1<sup>Flag</sup> | N/A | N/A |
| pBABE-TMEM186<sup>Flag</sup> | N/A | N/A |
| pLVX-COA1<sup>Flag</sup> | N/A | N/A |
| pLVX-COA1-GFP<sup>Flag</sup> | N/A | N/A |
| pLVX-COA1<sup>ΔIMS</sup>-GFP<sup>Flag</sup> | N/A | N/A |
| Chemicals and peptides |  |  |
| Anisomycin | Sigma-Aldrich | A9789 |
| EXPRE<sup>35</sup>S<sup>35</sup>S Protein Labeling Mix, [<sup>35</sup>S]-, 7mCi (74MBq), 50mM Tricine (pH 7.4), 10mM 2-mercaptoethanol | Perkin Elmer | NEG072007MC |
| Puromycin | InvivoGen | ant-pr-1 |
| Carbonyl cyanide 3'-chlorophenylhydrazone | Sigma-Aldrich | C2759 |
| Anti-Flag M2 Affinity gel | Sigma-Aldrich | A2220 |
| Flag peptide | Sigma-Aldrich | F3290 |
| Digitonin- High purity | Merck | 300410 |
| Dithiobis(succinimidyl propionate)<br>DSP | Peirce | 22585 |
| Streptavidin magnetic beads | ThermoFisher | 88816 |
| Biotin | Sigma-Aldrich | B4639 |
| Chloramphenicol | Sigma-Aldrich | C0378 |
| Proteinase K, from *Tritirachium album* | Sigma-Aldrich | P6556 |

### Cell lines and culturing

Cells were cultured in DMEM supplemented with 10% (v/v) fetal bovine serum (FBS), 1×penicillin/streptomycin (Sigma-Aldrich; P4458), 1×Glutamax™ (Life Technologies; 35050061) and 50µg/mL Uridine (Sigma-Aldrich; U3750). Cells were grown at 37°C with a 5% CO_2_ atmosphere. Galactose containing DMEM was prepared from Glucose-free DMEM (Life Technologies; 11966-025) supplemented with 10% (v/v) dialysed FBS, 25mM D-galactose (Sigma-Aldrich; G0750), 1×penicillin/streptomycin (Sigma-Aldrich; P4458), 1×Glutamax™ (Life Technologies; 35050-061), 1×Sodium Pyruvate (Life Technologies; 11360-070) and 50µg/mL Uridine (Sigma-Aldrich; U3750).

For the chloramphenicol and chase experiment, cells were grown for 4 days in glucose containing DMEM supplemented with 50µg/mL chloramphenicol (CAP) to inhibit mtDNA encoded protein synthesis and deplete respiratory chain complexes. Efficiency was verified by the lack of cell growth on galactose-DMEM. For the chase, cells were then plated onto 10cm plates and regular culture media was added. Cells were harvested at the desired time points. Mitochondria were then isolated and analysed by BN-PAGE.

For SILAC culture, cells were grown as previously described (Stroud et al., 2016). Briefly, cells were grown in DMEM without Lysine/Arginine (Assay Matrix; D9803-07B) supplemented with 10% (v/v) dialyzed FBS (GE Healthcare; SH30079.03), 1×penicillin/streptomycin (Sigma-Aldrich; P4458), 1×Glutamax™ (Life Technologies; 35050061), 1×Sodium Pyruvate (Life Technologies; 11360070) and 50µg/mL Uridine (Sigma-Aldrich; U3750) and either ‘heavy’ amino acids (^13^C_6_^15^N_4_-Arginine; Cambridge Isotope Labs Inc; CNLM-539-H-1, ^13^C_6_^15^N_2_-Lysine; Silantes; 211604102) or ‘light’ amino acids (Arginine; Sigma-Aldrich; A5131, Lysine; Sigma-Aldrich; L5626).

### Transfection and stable cell line generation

Cells were transfected using Lipofectamine LTX (Life Technologies; 15338100) according to manufacturer’s instructions. Stable, constitutive expression was achieved by transfecting HEK293T cells with pBABE-puro (Addgene; 1764) (Morgenstern and Land, 1990) encoding the cDNA of interest. Stable, inducible expression was achieved by transfecting cells with pLVX-TetOne-Puro (Clonetech; 631849) encoding the cDNA of interest. All plasmids were sequence verified by Micromon (Monash University) or the Garvin Institute for Medical Research. Viral supernatants were collected after 48 hours and used to infect the corresponding KO cell lines. Cell lines were then selected by growth on galactose-containing DMEM, with the exception of TMEM186 and COA1 knockouts, which were selected for by the addition of 2µg/mL puromycin. Expression was verified by BN-PAGE or SDS-PAGE as required.

### Gene editing and Screening

Gene editing was performed using TALEN pairs as described (Formosa et al., 2015) or the pSp-CAS9(BB)-2A-GFP (PX458) CRISPR/Cas9 construct (a gift from F. Zhang; Addgene plasmid 48138 (Ran et al., 2013)). TALEN pairs were designed using ZiFiT Targeter (Sander et al., 2010) and CRISPR/Cas9 guide RNAs were designed using CHOPCHOP (Montague et al., 2014). KO cell lines for NDUFAF1 and TIMMDC1 have been previously reported (Lake et al., 2019; Stroud et al., 2016). Target sequences, gene disruption strategies and generated indels can be found in Supplementary Table 1.

### Mitochondrial isolation, gel electrophoresis, immunoblot analysis and antibodies

Mitochondria were isolated as previously described (Johnston et al., 2002). Protein concertation was determined by bicinchoninic acid assay (BCA; Thermo Fisher Scientific; 23223). Mitochondria were used immediately or aliquoted and frozen at −80°C until required. Alkali extraction and protease protection assay was performed as previously described (Formosa et al., 2015). Tris-Tricine SDS-PAGE and BN-PAGE were performed as previously described (McKenzie et al., 2007; Schagger and von Jagow, 1987; Wittig et al., 2006). Membrane association and protease protection assay were performed as previously described (Ryan et al., 2001). Immunoblotting onto PVDF membrane (Merck; IPVH00010) was performed using a Novex Semi-Dry Blotter (Thermo Fisher) according to manufacturer’s instructions. Primary antibodies, sources and associated dilutions are listed above. Anti-mouse (Sigma-Aldrich; A9044) or anti-rabbit (Sigma-Aldrich; A0545) horseradish peroxidase-conjugated secondary antibodies were used at 1 dilution of 1:10,000. Clarity western ECL chemiluminescent substrate (BioRad; 1705061) was used for detection on a BioRad ChemiDoc XRS+ imagining system.

### Proximity biotin identification (BioID)

Biotin labelling and identification was performed as previously described with some modifications (Osellame et al., 2016; Roux et al., 2012). Briefly, NDUFAF1^KO^ cells expressing NDUFAF1^BirA*^ or NDUFAF1^HA^ were grown in standard DMEM supplemented with 50µM Biotin for 24 hours. Isolated mitochondria were solubilized in RIPA buffer (50mM Tris-HCl pH 7.5, 150mM NaCl, 1% NP-40, 1mM EDTA, 1mM EGTA and 0.1% SDS), supplemented with 1× protease cocktail inhibitor (Sigma-Aldrich; 11873580001), 0.5% sodium deoxycholate and 250U/mL benzonase (Sigma-Aldrich; E1014) and biotinylated proteins were enriched using streptavidin-coated magnetic beads (ThermoFischer; 88816). On-bead tryptic digest was performed using 5µg/mL mass spectrometry grade trypsin (Promega; V5113) overnight at 37°C with agitation. The beads were rinsed twice with mass spectrometry grade water, the rinse fraction was pooled with tryptic digests and peptides were then concentrated using vacuum centrifugation. The resulting pellets were resuspended in 0.1% TFA, 2% ACN prior to analysis by LC-coupled mass-spectrometry.

### Affinity enrichment

For affinity enrichment of Flag-tagged proteins, mitochondria from control or Flag-expressing cells were isolated and solubilized in 1% (w/v) digitonin or 1% (v/v) Triton X-100 in solubilization buffer (20mM Bis-Tris (pH 7.0), 50mM NaCl, 10% (v/v) glycerol). Following clarification of lysates by centrifugation (16,000*g*, 10 mins, 4°C), supernatants were applied to Flag affinity gel (Sigma) and incubated for 2 hours at 4°C with gentle rotation. Affinity gel was washed with 0.1% digitonin or 0.1% Triton X-100 in solubilization buffer as required. Proteins were eluted using 150 µg/mL Flag peptide in solubilization buffer with 0.1% (w/v) digitonin or 0.1% (v/v) Triton X-100 as required. Samples were then prepared for BN-PAGE or proteomics analysis as described.

### mtDNA- encoded protein metabolic labelling and *in vitro* protein import

Radiolabelling of mtDNA-encoded proteins was performed using 10µg/mL Anisomycin (Sigma-Aldrich; A9789) to block cytosolic translation as previously described (Formosa et al., 2016). Isolated mitochondria were subjected to Flag affinity enrichment and/or SDS-PAGE analysis, followed by transfer to PVDF membrane and analysed by phosphorimaging digital autoradiography (GE Healthcare).

For affinity enrichment of newly translated proteins, mitochondria were isolated after 2 hours of labelling with [^35^S]-Methionine/Cysteine mix and affinity enrichment performed as above. Bound proteins were removed from beads by incubating affinity gel in LDS sample buffer (Thermo Fisher; NP0008) with 100mM DTT at room temperature for 15 minutes. Affinity gel was removed by centrifugation and proteins analysed by Tris-Tricine SDS-PAGE as described above.

For co-immunoprecipitation of endogenous NDUFAF1 or ECSIT, 143B TK^-^ osteosarcoma cells were labelled for 2 hours with [^35^S]-Methionine/Cysteine mix and chased for 0, 3 and 24 hours in DMEM. Isolated mitochondria were solubilized in 1% Triton X-100 and incubated with Protein A sepharose crosslinked to pre-immune sera or NDUFAF1/ECSIT reactive sera. To determine direct protein-protein interactions, samples were initially incubated with 0.2mM DSP in SM buffer (250mM sucrose, 10mM MOPS pH 7.2) followed by solubilization in 1% Triton X-100 and 1% SDS. Protein-antibody complexes were washed, and bound proteins were eluted in 100mM Glycine pH 2.0 and prepared for SDS-PAGE with or without 100mM DTT as indicated.

For *in vitro* protein import, the cDNA sequence of TMEM186 was amplified to incorporate a T7 promoter sequence (5’-GTACCGTAATACGACTCACTATAG-3’) upstream of the initiation codon and a reverse primer to incorporate a poly-A tail (5’-T_18_CACTTGAGCATCTGATGTACCC-3’). Radiolabelled protein was generated using the TNT-T7 quick-coupled transcription/translation system (Promega; L1170). Translated protein was incubated at 37°C with isolated mitochondria as previously described (Lazarou et al., 2007). Following import, mitochondria were prepared for and analysed by SDS-PAGE and phosphorimaging.

### Proteomics analysis

Stable isotope labelling of amino acids in cell culture (SILAC) was performed as described (Stroud et al., 2016). BioID proteomics was performed as previously described (Osellame et al., 2016). Affinity enrichment proteomic analysis was performed as previously described (Thompson et al., 2018).

#### Mass spectrometry of SILAC treated cells

Mitochondria were isolated as above from COA1^KO^ cells cultured in SILAC DMEM and prepared as described previously (Stroud et al., 2016). Analysis was performed on SILAC labelled whole cell lysates of the COA1^KO^ cell line. Mitochondrial protein yield and whole cell pellet protein amounts were assayed by BCA. For each triplicate set, two replicates contained 50 μg of protein from heavy labelled cells/mitochondria were mixed with the same amount of protein from light KO cells/mitochondria. The third replicate consisted of a label switch. Both whole cell and mitochondrial samples were treated as follows: Samples were solubilized in 1% (w/v) SDC, 100mM Tris-Cl pH 8.1, 40mM chloroacetamide and 10mM TCEP prior to vortexing and heating for 5 minutes at 99 °C with 1500 rpm shaking. Samples were then sonicated for 15 minutes in a room temperature water bath prior to the addition of 1 μg trypsin (Promega) and incubation overnight at 37 °C. The supernatant was then transferred to 3x 14G 3M™Empore™ SDB-RPS stage tips (Kulak et al., 2014). Ethyl acetate (99%) and 1% TFA was added to the tip before centrifugation at 3000*g* at room temperature as described (Kulak et al., 2014). Stage tips were washed first with 99% ethyl acetate and 1% TFA and then with ethyl acetate supplemented with 0.2% TFA. Samples were eluted in 80% acetonitrile (ACN) and 1% NH_4_OH, and acidified to a final concentration of 1% TFA prior to drying in a SpeedVac.

#### Mass-spectrometry and data analysis

Peptides were reconstituted in 2% ACN, 0.1% TFA and transferred to autosampler vials for analysis by online nano-HPLC/electrospray ionization-MS/MS on either a Thermo-Fisher Scientific Orbitrap Q Exactive Plus, Orbitrap Fusion Lumos Tribrid or Orbitrap Elite Hybrid Ion-Trap instrument as follows:

NDUFAF1^BirA*^, TMEM186^Flag^ and COA1^Flag^ AE-MS were acquired on an Orbitrap Elite Hybrid Ion-Trap instrument connected to an Ultimate 3000 HPLC (Thermo-Fisher Scientific). Peptides were loaded onto a trap column (PepMap C18 trap column 75 µm x 2 cm, 3 µm, particle size, 100 Å pore size; ThermoFisher Scientific) at 5 μL/min for 3 min before switching the pre-column in line with the analytical column (PepMap C18 analytical column 75 μm x 50 cm, 2 μm particle size, 100 Å pore size; ThermoFisher Scientific). Peptide separation was performed at 300 nL/min using a 90 min non-linear ACN gradient of buffer A [0.1% (v/v) formic acid, 2% (v/v) ACN, 5% DMSO] and buffer B [0.1% (v/v) formic acid in ACN, 5% DMSO]. Data was collected in Data Dependent Acquisition (DDA) mode using m/z 300 - 1650 as MS scan range, rCID for MS/MS of the 20 most intense ions. Other instrument parameters were: MS scan at 120,000 resolution, maximum injection time 150 ms, AGC target 1E6, CID at 30% energy for a maximum injection time of 150 ms with AGC target of 5,000.

For COA1^KO^ mitochondria, data were acquired on an Orbitrap Fusion Lumos instrument connected to an Ultimate 3000 HPLC. Peptides were loaded onto a trap column (PepMap C18 trap column 75 µm x 2 cm, 3 µm, particle size, 100 Å pore size; ThermoFisher Scientific) at 5 μL/min for 3 min before switching the pre-column in line with the analytical column (PepMap C18 analytical column 75 μm x 50 cm, 2 μm particle size, 100 Å pore size; ThermoFisher Scientific). The separation of peptides for was performed at 300 nL/min using a 90 min non-linear ACN gradient of buffer A [0.1% (v/v) formic acid, 2% (v/v) ACN, 5% DMSO] and buffer B [0.1% (v/v) formic acid in ACN, 5% DMSO]. The mass spectrometer was operated in positive-ionization mode with spray voltage set at 1.9 kV and source temperature at 275°C. Lockmass of 401.92272 from DMSO was used. Data were collected using the Data Dependent Acquisition using m/z 350-1550 at 120000 resolution with AGC target of 5e5. The “top speed” acquisition method mode (3 sec cycle time) on the most intense precursor was used whereby peptide ions with charge states ≥2-5 were isolated with isolation window of 1.6 m/z and fragmented with high energy collision (HCD) mode with stepped collision energy of 30 ±5%. Fragment ion spectra were acquired in Orbitrap at 15000 resolution. Dynamic exclusion was activated for 30s.

Raw files were analysed using the MaxQuant platform (Tyanova et al., 2016a) version 1.6.5.0 searching against the UniProt human database containing reviewed, canonical entries (January 2019) and a database containing common contaminants. For label-free (LFQ) AE-MS experiments, default search parameters were used with “Label free quantitation” set to “LFQ” and “Match between runs” enabled. Using the Perseus platform (Tyanova et al., 2016b) version 1.6.7.0, proteins group LFQ intensities were Log_2_ transformed. Values listed as being ‘Only identified by site’, ‘Reverse’ or ‘Contaminants’ were removed from the dataset. Mitochondrial annotations were imported by matching with the Mitocarta2 dataset (Calvo et al., 2016) by gene name and/or ENSG identifier and the Integrated Mitochondria Protein Index (IMPI) database by gene name (Smith and Robinson, 2019). Experimental groups were assigned to each set of triplicates and the number of valid values for row group calculated. For each experiment (containing a control and an enrichment group) rows having less than 3 valid values in the enrichment group were removed and the missing values in the relevant control group imputed to values consistent with the limit of detection. A modified two-sided Student’s t-test based on permutation-based FDR statistics (Tyanova et al., 2016b) was performed between the two groups. The negative logarithmic *p*-values were plotted against the differences between the Log_2_ means for the two groups. The significance threshold used for these experiments is noted in the relevant figure legend and supplementary tables. For SILAC experiments, default search parameters were used with multiplicity set to 2 (Lys8, Arg10) and “Match between runs” enabled. Using the Perseus platform (Tyanova et al., 2016b) version 1.6.7.0, proteins group normalized H/L ratios were Log_2_ transformed. Label switched samples (L/H) were inverted to KO/HEK293T orientation prior to this step. Values listed as being ‘Only identified by site’, ‘Reverse’ or ‘Contaminants’ were removed from the dataset. Mitochondrial annotations were imported by matching with the Mitocarta2 and IMPI datasets as above. Experimental groups were assigned to each set of triplicates rows with <2 valid values for each group removed. Non-mitochondrial proteins were removed, and a one sample Student’s two-sided t-test was performed within each group. The negative logarithmic *p*-values were plotted against the differences between the mean ratios for each group. A significance threshold (*p* < 0.05) was used for all experiments. Log_2_-transformed median SILAC ratios were mapped on homologous subunits of the respiratory chain complexes. Applicable Protein Data Bank accession codes provided in text.

#### Experimental Design and Statistical Rationale

SILAC labelled whole cell and mitochondrial analyses were performed in triplicate, with a label switch as we have done previously (Dibley et al., 2017; Stroud et al., 2015; Stroud et al., 2016). The statistical approaches used to analyse the data was consistent with published quantitative SILAC analyses from our (Stroud et al., 2015; Stroud et al., 2016) and other labs employing similar instrumentation and methods. For steady state analysis the fold change threshold we used for significance was >1.5 and a *p*<0.05 following a single sample two-sided Student’s t-test. AE-MS analyses were performed in triplicate using label free quantitation and compared to untagged control cells as we (Lim et al., 2018) and others have done previously using similar instrumentation and methods. Imputation was performed only on missing values in control samples and random values were drawn from a narrow distribution equivalent to the limit of detection of the instrument. Significantly-enriched proteins were determined through a modified two-sample two-sided Student’s t-test based on permutation-based FDR statistics (Tyanova et al., 2016b) between the two groups, with an FDR of 1% and the s0 value determined iteratively to exclude any enriched proteins specific to the control group.

### Image processing and quantitation

Quantified data was measured using ImageLab (BioRad) or ImageJ (Schneider et al., 2012) and was then prepared in GraphPad Prism 7. All figures were prepared using Adobe Photoshop/Illustrator (CC2019).

