## Supplementary Figures for "Dissecting the Roles of Mitochondrial Complex I Intermediate Assembly (MCIA) Complex Factors in the Biogenesis of Complex I"

### Supplemental Figures S1-S4

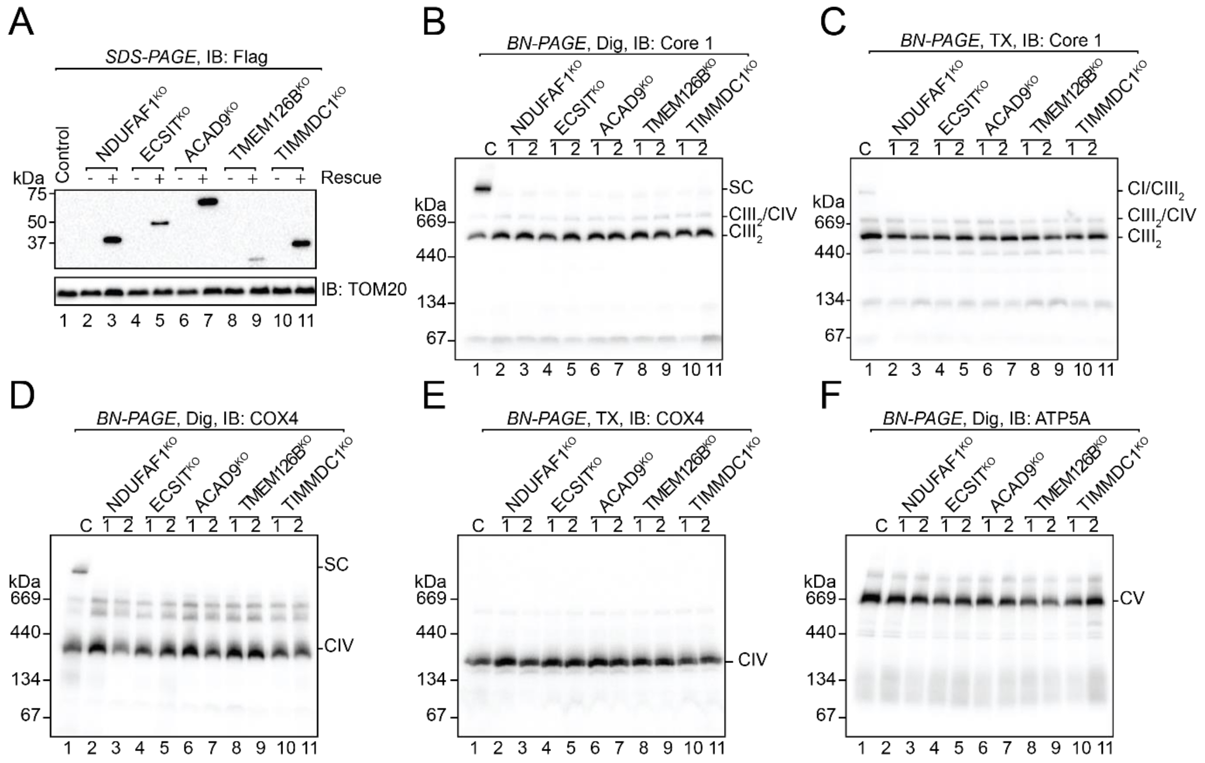


Formosa et al., Supp. Figure 1

**Supplementary Figure 1 (related to Figure 2): Loss of the MCIA complex and TIMMDC1 results in a specific defect in complex I. (A)** Mitochondrial proteins from control, knockout and rescued cell lines were separated by SDS-PAGE and subjected to western blot analysis using Flag antibodies. TOM20 served as a loading control. Mitochondria from control and knockout cells were subjected to BN-PAGE and immunoblotting as follows **(B)** digitonin, Core 1 (CIII), **(C)** Triton X-100, Core 1 (CIII), **(D)** digitonin, COX4 (CIV), **(E)** Triton X-100, COX4 (CIV) and **(F)** digitonin, ATP5A (CV).

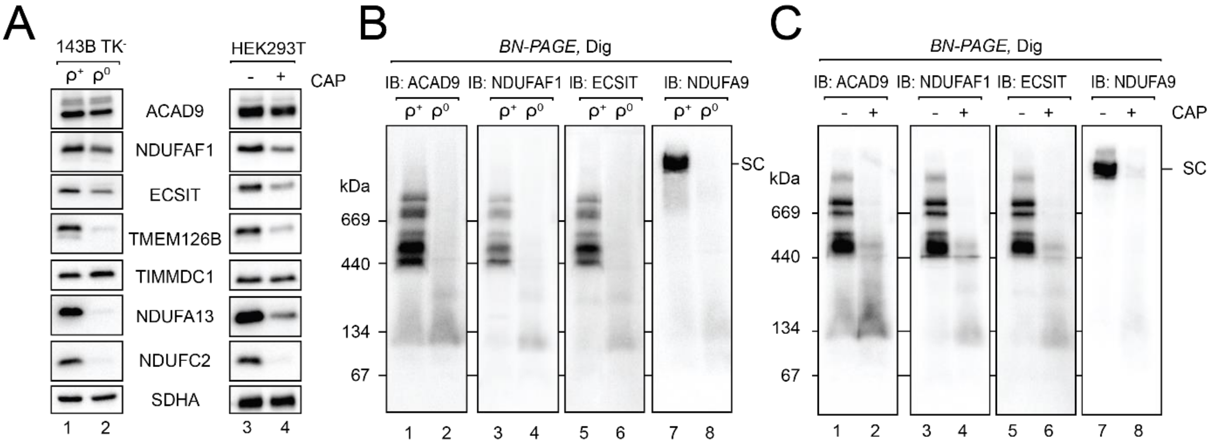


Formosa et al., Supp Figure 2

**Supplementary Figure 2 (related to Figure 3): Stability of the MCIA complex is coupled to translation (A)** Mitochondrial proteins isolated from 143B TK$^-$ wildtype ($\rho^+$) or depleted of mtDNA ($\rho^0$) or control HEK293T cells grown with or without chloramphenicol (CAP) were subjected to SDS-PAGE and immunoblotting using antibodies as indicated. SDHA was used as a loading control. **(B)** Mitochondria isolated from 143B TK$^-$ $\rho^+$ or $\rho^0$ were subjected to BN-PAGE and immunoblotting using antibodies as indicated. **(C)** Mitochondria isolated from HEK293T cells grown in the with or without chloramphenicol were subjected to BN-PAGE and immunoblotting using antibodies as indicated.

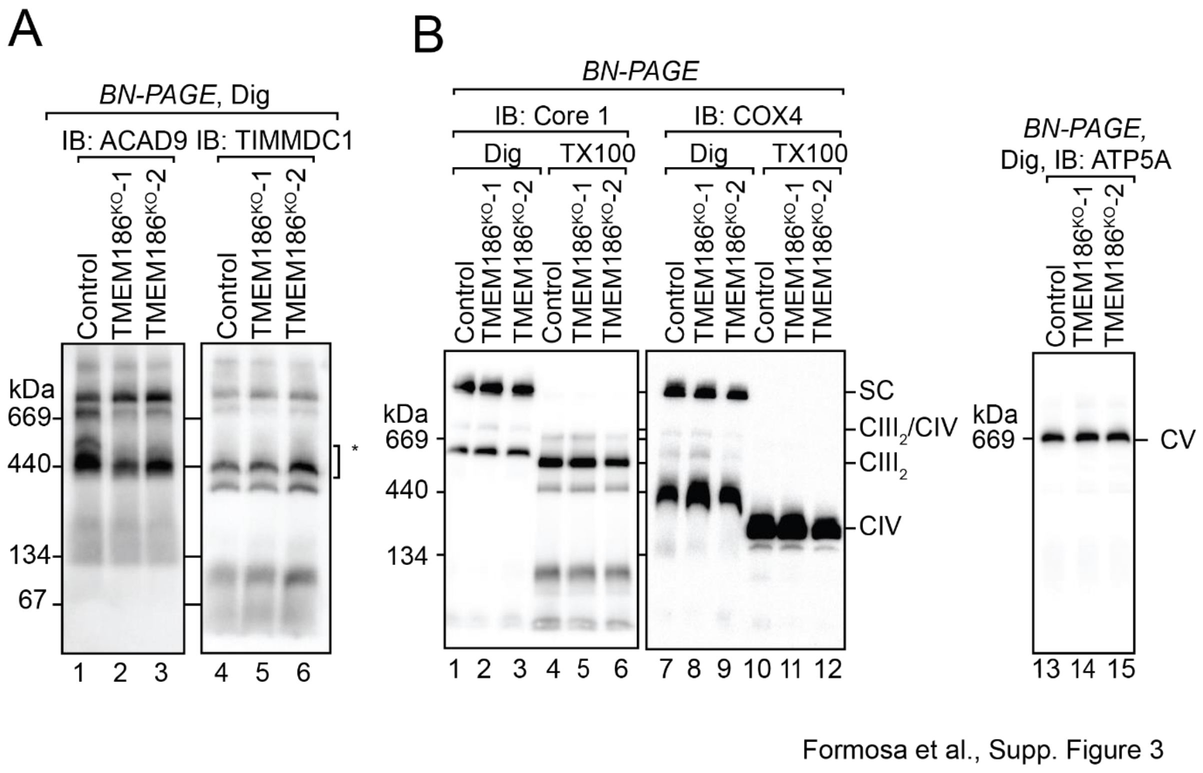


**Supplementary Figure 3 (related to Figure 4): Analysis of complex I assembly intermediates and OXPHOS complexes in TMEM186$^{KO}$ mitochondria (A)** Isolated mitochondria from control and TMEM186$^{KO}$ cells were solubilized in 1% digitonin and subjected BN-PAGE and immunoblotting with ACAD9 or TIMMDC1 antibodies. **(B)** Isolated mitochondria from control and TMEM186$^{KO}$ cells were solubilized in 1% digitonin or 1% Triton X-100 and subjected BN-PAGE and immunoblotting with Core 1 (CIII), COX4 (CIV) or ATP5A (CV) antibodies.

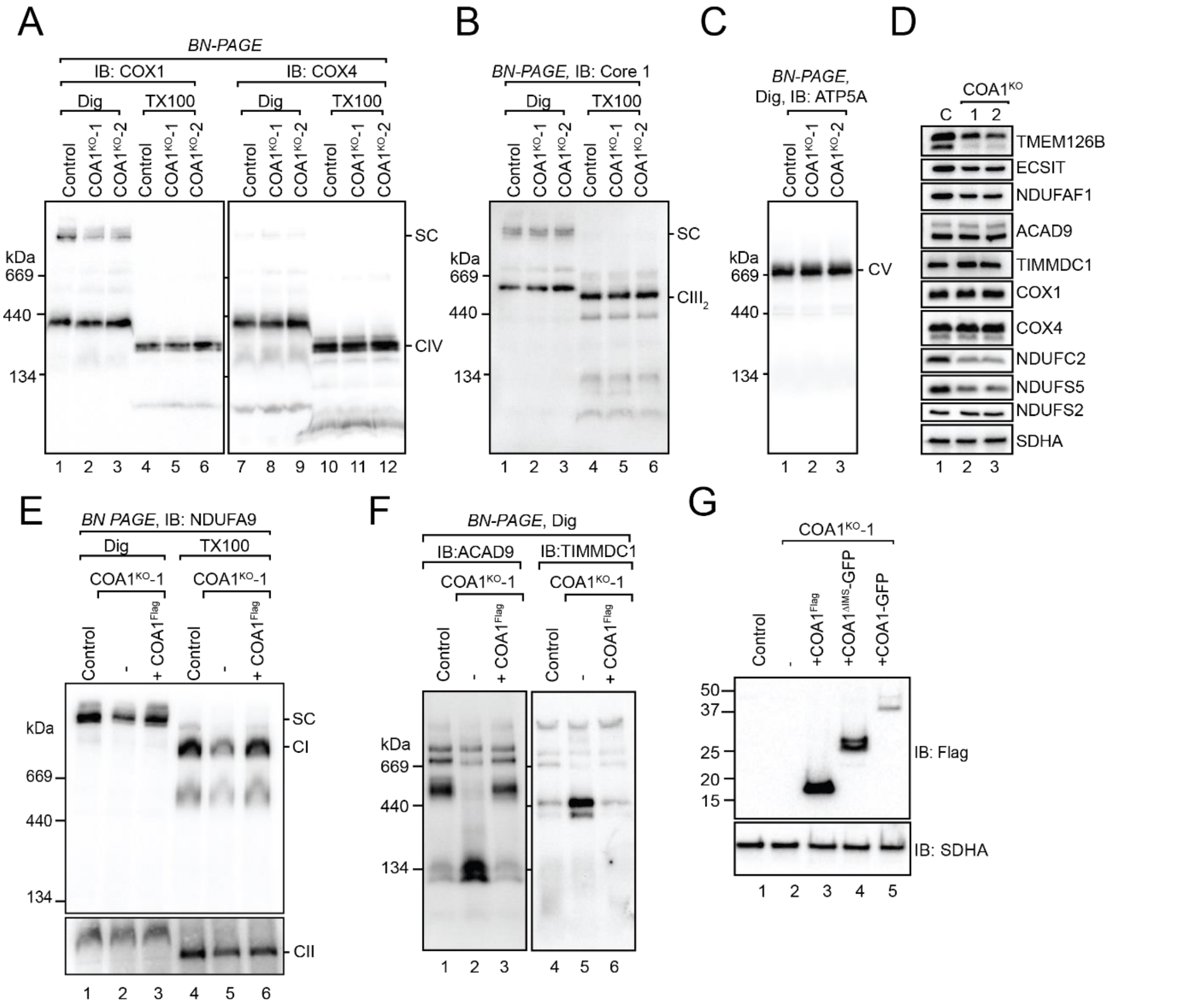


Formosa et al., Supp. Figure 4

**Supplementary Figure 4 (related to Figure 5): COA1 is required for complex I assembly but dispensable for complex IV assembly (A)** Isolated mitochondria from control and $COA1^{KO}$ cells were solubilized in 1% digitonin or 1% Triton X-100 and subjected BN-PAGE and immunoblotting with COX1 or COX4 antibodies. **(B)** Isolated mitochondria from control and $COA1^{KO}$ cells were solubilized in 1% digitonin or 1% Triton X-100 and subjected BN-PAGE and immunoblotting with Core 1 antibodies. **(C)** Isolated mitochondria from control and $COA1^{KO}$ cells were solubilized in 1% digitonin and subjected BN-PAGE and immunoblotting with ATP5A antibodies. **(D)** Isolated mitochondria from control and $COA1^{KO}$ cells were analysed by SDS-PAGE and immunoblotting with antibodies as indicated. **(E)** Mitochondria from control, $COA1^{KO}$ or $COA1^{KO}$ + $COA1^{Flag}$ were isolated, solubilized in 1% digitonin or 1% Triton X-100 and subjected to BN-PAGE. Immunoblotting was performed with antibodies directed to NDUFA9. **(F)** Mitochondria from control, $COA1^{KO}$ or $COA1^{KO}$ + $COA1^{Flag}$ were isolated and subjected to BN-PAGE analysis after solubilization in 1% digitonin. Immunoblotting was performed with antibodies directed to ACAD9 and TIMMDC1. **(G)** Mitochondria from control, $COA1^{KO}$ and $COA1^{KO}$ cells expressing $COA1^{Flag}$, $COA1\text{-}GFP^{Flag}$ and $COA1^{\Delta IMS}\text{-}GFP^{Flag}$ were subjected to SDS-PAGE and immunoblotting with Flag antibodies. SDHA was used as a loading control.
